# Dynamic polyadenylation safeguards developmental trajectories during paused embryogenesis

**DOI:** 10.64898/2026.05.18.726068

**Authors:** Christopher Phillip Chen, Hannah Greenfeld, Sarah Foust, Daniel Wagner

## Abstract

Embryonic pausing, including forms of diapause, enables development to be reversibly suspended during adverse conditions, but how paused embryos preserve cell fate patterning remains unclear. Using zebrafish, we show that developmental pausing results in the temporary collapse of Wnt, BMP, FGF, and Nodal signaling gradients, despite broad preservation of cell identity at the transcriptional level. In parallel, pausing induces a dormancy-enriched gene expression program (DEEP), which includes the non-canonical poly(A) polymerase *tent5ba.* Tent5ba promotes survival through pausing, sustains DEEP expression, and is required for robust reestablishment of axial patterning upon developmental re-entry. Poly(A)-tail and transcriptional profiling further link *tent5ba* to the stabilization of mRNA targets associated with robust developmental outcomes, supporting a model in which transcript polyadenylation safeguards patterning fidelity through suspended embryogenesis.

## Main Text

Embryos of all species, including humans, endure persistent intrinsic and extrinsic challenges, yet generally succeed in achieving stereotyped developmental outcomes(*1*, *2*). Reversible pausing of embryogenesis (e.g., diapause) represents an evolutionarily widespread resilience strategy in which developmental progression is coupled to the presence of favorable environmental or metabolic conditions(*3*, *4*). In zebrafish embryos, for example, rapid and reversible developmental arrest can be triggered by hypoxia, metabolic inhibition, and/or physical crowding (Fig. SlA) (*5–8*). Although embryonic pausing is well documented(*4*), the underlying molecular mechanisms - particularly those that sustain developmental patterning - remain unknown. Here, we leverage embryonic zebrafish as a scalable model to induce, molecularly profile, and genetically perturb the process of developmental pausing. Our findings reveal a striking decoupling during paused embryogenesis: morphogen signaling gradients collapse, while cell fates are largely preserved at the transcriptional level. In addition, paused embryos simultaneously upregulate a protective <u>“Dormancy-Enriched</u> gene ;Expression £rogram” (DEEP) that promotes pausing tolerance and axial patterning fidelity. Guided by genetic analysis of a key DEEP factor, *tent5ba,* a non-canonical poly(A) polymerase, we identify poly(A) tail regulation as a previously unappreciated post-transcriptional mechanism that preserves transcriptional trajectories through pausing, supporting successful developmental resumption.

### Embryonic pausing leads to the collapse of morphogen signaling

To explore the molecular attributes of morphogen-based axis patterning during pausing, we implemented and extended established methods (7, *8*), for hypoxia- or potassium cyanide-(KCN) based arrest of embryogenesis (Fig. lA). Quantifications of epiboly progression, a stage-defining feature of gastrulation(9), revealed that embryos exposed to hypoxia experience rapid-onset developmental pausing (Fig. lB). Following reoxygenation, the majority of paused embryos resumed normal development, survived through larval stages (Fig. SlB-D) and could grow to become sexually viable adults (Fig. SlE). While the majority of reoxygenated embryos appeared phenotypically normal, a rare but consistent proportion (∼1.3%) developed mild phenotypes consistent with axis patterning disruption, e.g. loss of ventral tail fin structures (Fig. SlF-G)(*10*). Taken together, these data confirmed that zebrafish embryos can reversibly pause developmental progression with minimal consequences for viability and developmental patterning.

Upon reoxygenation, paused embryos resume development from an early gastrula stage after 18 hours (or more) of arrest, a time interval that would normally span gastrulation, convergent extension, segmentation, and the early stages of organogenesis. This temporal distortion requires that paused embryos preserve key attributes of early gastrulation over unusually prolonged intervals. To test whether paused embryos retained molecular features of gastrula-stage axial patterning, we assayed the activity of the central pathways that mediate morphogen signaling during gastrulation: canonical Wnt, Bone Morphogenic Protein (BMP), Nodal, and Fibroblast Growth Factor (FGF)(*11–14*). In control embryos, we observed characteristic signaling activity gradients consistent with previous studies (Fig. 1C-H, S2A-H)(*11–14*). By contrast, both the intensity and shape of Wnt, BMP, FGF, and Nodal signaling gradients were effectively abolished in paused embryos (Fig. lC-H, S2A-H). This striking result revealed that paused embryos, which resume normal development with high fidelity, successfully endure dynamic erasure of developmental signaling states. It also suggested that additional dimensions of identity (beyond signaling) might be critical for maintaining state identity during periods of extended embryonic pausing.

**Fig. 1.**
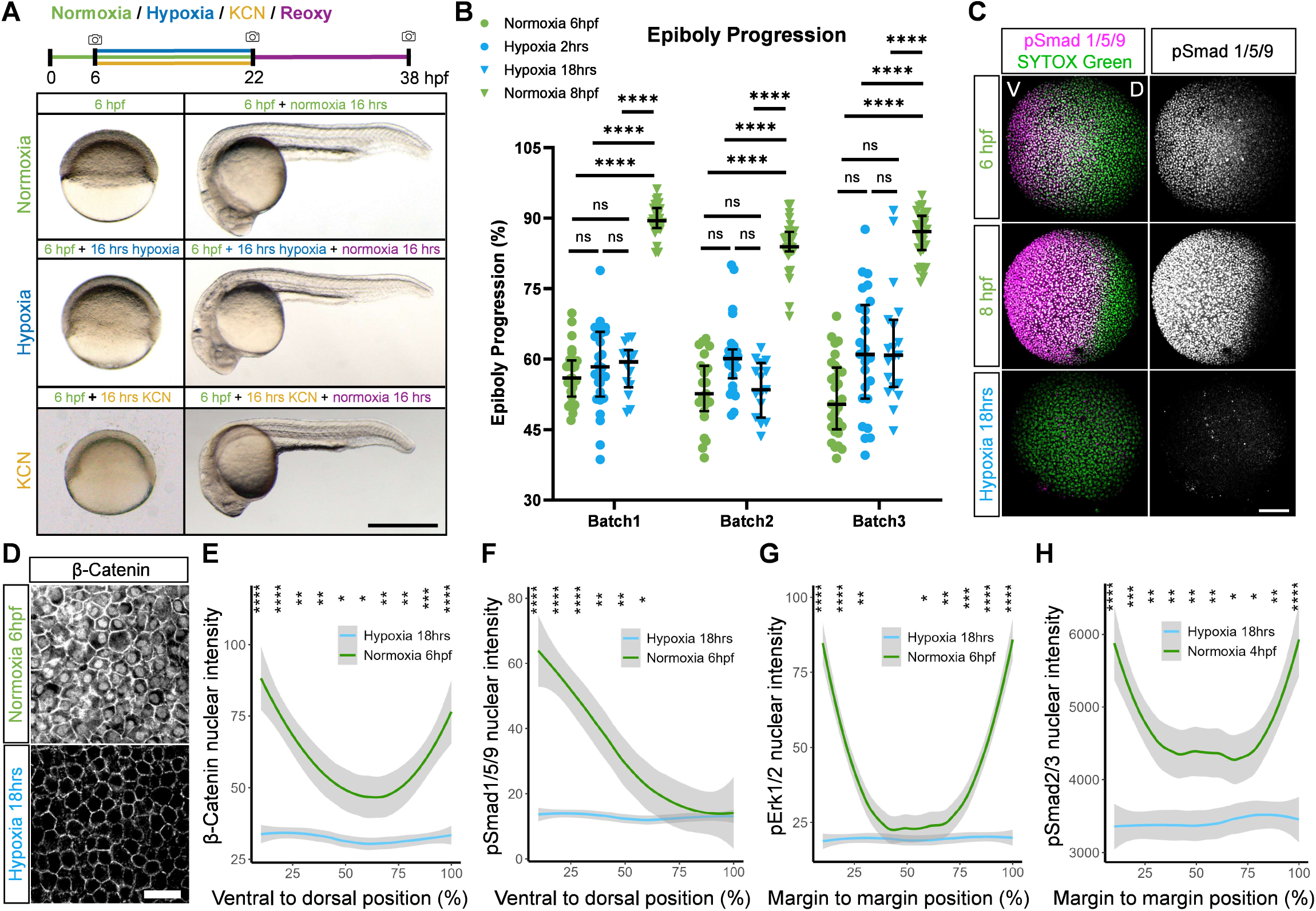
Reversible embryonic pausing and loss of morphogen signaling activity in response to hypoxia. **(A)** Left, representative brightfield images of shield-stage normoxic control embryos (green) and shield state embryos exposed to 16 hrs hypoxia (blue) or KCN (yellow). Right, embryos imaged after an additional 16 hrs in normoxia or reoxygenation recovery. Scale, 578 µm. **(B)** Epiboly progression (%), median with interquartile range, from three separate experiments. 6 hpf embryos were fixed immediately or after 2 hrs hypoxia, 18 hrs of hypoxia, or 2 hrs of normoxia. In all figures, (P or adjusted P-values): * = P≤0.05, ** = P≤0.01, *** = P≤0.001, **** = P≤0.0001 (Methods). Kruskal-Wallis tests. **(C, D)** Representative whole-mount immunofluorescence (IF) images. (C) IF pSmadl/5/9 and SYTOX Green, maximum intensity projections (200 µm). (D) IF β-Catenin, single confocal plane, shield stage margin. See Fig. S2A. Scale bars (C) 26 µm, (D) 120 µm. **(E-H)** Quantifications of β-Catenin, pSmadl/5/9, pErkl/2, or pSmad2/3 nuclear intensity (E-G) normoxic 6 hpf and 6 hpf after 18 hrs hypoxia or (H) normoxic 4 hpf and 4 hpf after 18 hrs hypoxia. Solid lines indicate smoothed conditional means of embryos, 95% confidence interval in gray. See Fig. S2. Wilcoxon test across deciles.

### Paused embryos maintain cellular identities at the transcriptional level

To determine whether cells in paused embryos maintain their identities at the transcriptional level, we performed single-cell RNA-sequencing (scRNA-seq) on dissociated embryo cell suspensions collected during normal, paused, and reoxygenated development (Fig. 2A). The resulting datasets were processed, and annotated by cell type label transfer from ZMAP (*15*), a multi-study consensus scRNA-seq atlas of zebrafish development (Fig. 2B, Methods). Co-clustering of perturbed scRNA-seq profiles over both assigned cell type labels and the harmonized UMAP embedding indicated that cells from paused embryos resembled those of 6 hpf control embryos, while cells from reoxygenated embryos resembled control cells from 6 and/or 8 hpf (Fig. 2B-C). To further quantify transcriptional differences, we computed Euclidean distances in principal component (PC) space between cell group centroid pairs annotated by both cell type labels and perturbation condition (Fig. 2D, S3A). Centroid pairs sharing the same cell type labels from different conditions were consistently more similar (displayed shorter distances) than within-condition pairs between cell types (Fig. 2D, S3A-B). Cell types also displayed consistent expression of key developmental marker genes regardless of condition (Fig. 2E, S3C). Together, these data indicated that cells of both paused and recovering embryos retained stereotypic patterns of cell-type-specific gene expression seen during normal development.

**Fig. 2.**
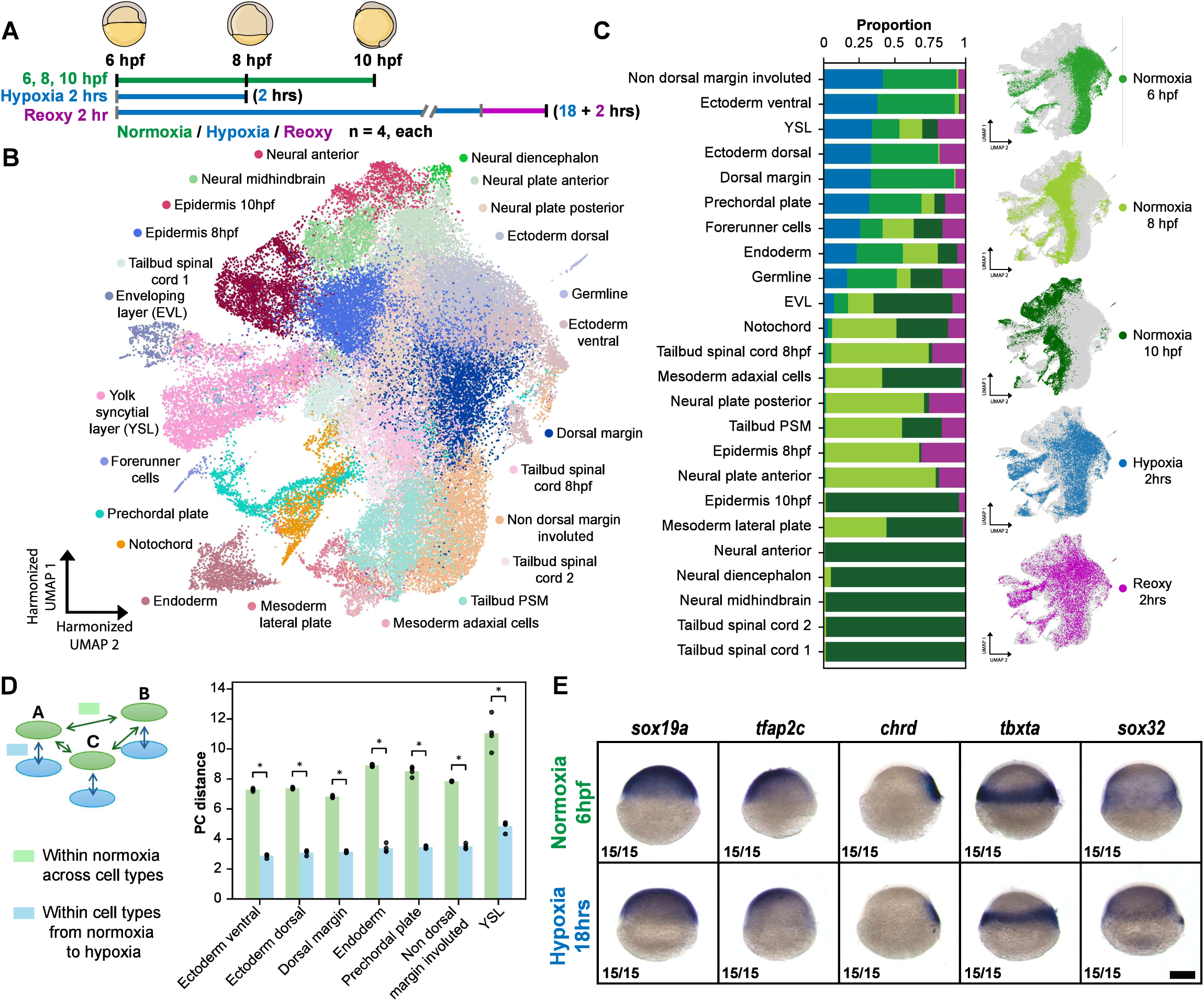
Preservation of cell-type specific transcriptomes during paused embryogenesis. **(A)** Summary of scRNA-seq time-points and conditions: normal development (6, 8, and 10 hpf), paused development (6 hpf + 2 hrs of hypoxia), and reoxygenated development (6 hpf + 18 hrs hypoxia+ 2 hrs of reoxygenation). **(B)** Harmonized UMAP embedding colored by cell type annotations. **(C)** Left, cell type compositions by sample group, ordered by hypoxia proportion. Right, UMAP insets colored by sample groups. **(D)** Comparison of Euclidean distance in principal component space between cell type-sample group pairs. Mean distances spanning between control cell types (green) and spanning conditions within cell types (blue), are plotted for the 7 most abundant cell types. Mann-Whitney U test (n=4). **(G)** Whole-mount in situ hybridization for *soxl 9a, tfap2c, chrd, tbxta,* and *sox32* in embryos at 6 hpf or 6 hpf after 18 hrs of hypoxia. Animal pole up, shield/dorsal right and ventral left. Images represent n=15 over 3 biological replicates. Scale, 200 µm. In all figures, (P or adjusted P-values): * = P≤0.05, ** = P≤0.01, *** = P≤0.001, **** = P≤0.0001 (Methods).

### Paused embryos express a <u>Dormancy-Enriched Expression</u> E,rogram (DEEP)

Alongside the retention of cell-type specific transcriptional profiles, paused embryos activated novel transcriptional programs not typically observed during normal development. These changes were revealed through the application of consensus non-negative matrix factorization (cNMF)(*16*), which identified dozens of gene expression programs (GEPs) corresponding to stable cell types and dynamic states (Fig. S4A-B, Data Sl). Two such dynamic programs, GEP5 and GEP12, were elevated in all major cell type clusters in paused embryos and during reoxygenated recovery, respectively (Fig. S4C-D). These trends were confirmed by differential expression analysis, pseudobulked across cell type clusters, which revealed 52.1% of differentially expressed genes in paused conditions (log2FC>0.5, P-adj<0.05) were associated with GEP5 (Fig. S4E, Data S2). During developmental pausing, embryos thus preserve cell-type specific transcriptional identities while simultaneously upregulating a coordinated global gene expression response, which we refer to herein as a <u>“Dormancy-Enriched</u> gene _Expression £rogram” (DEEP).

To further investigate DEEP dynamics and composition, we performed bulk RNA-seq and bulk Assay for Transposase-Accessible Chromatin sequencing (ATAC) with control, paused, and recovered embryos at multiple timepoints (Fig. 3A, SSA). Principal component analysis of global gene expression patterns revealed orthogonal relationships between embryonic development and rapid, reversible pausing dynamics (Fig. 3A, S5A-B). Differential gene expression (DGE) comparisons between prolonged paused and control embryos identified 2493 differentially expressed genes (DEGs, P-adj<0.01, llog2FCl>l) (Fig. 3B, SSC, Data S3), while Gene Set Enrichment Analysis (GSEA) revealed an enrichment of key pathways amongst DEEP genes: glycolysis/gluconeogenesis *(ldha, gapdhs, enol a),* FoxO/insulin signaling *(foxol a, irs2a, gad45bb),* cellular senescence *(cdknld, serpinel, atm),* Wnt signaling *(wntl, wnt2, myca),* MAPK signaling (*egfra, fgfl 6, fosab*), and a suppression of aminoacyl-tRNA biosynthesis *(sepsecs, qrsll, dars2),* many of which are hallmarks of dormancy programs in other species (Fig. 3C, Data S5)(3, *17–19*). Differential accessibility analysis confirmed that a majority (67%) of upregulated transcripts in prolonged hypoxia compared to normal development were proximal ([-2000, +500] bp from TSS) to a peak that increased in accessibility (Fig. S5D, Data S4).

**Fig. 3.**
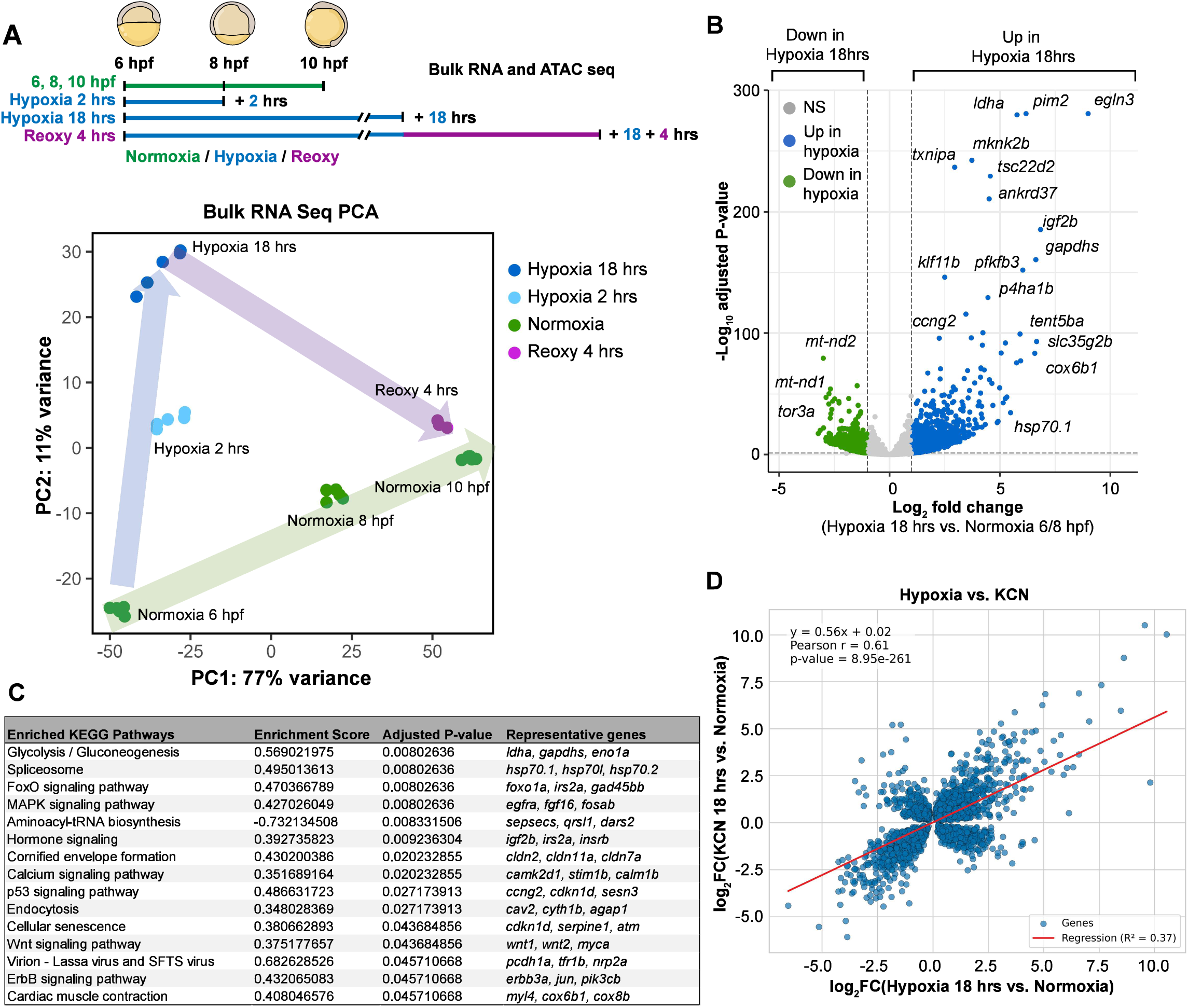
Expression dynamics of the Dormancy Enriched Expression Program (DEEP) **(A)** Summary of bulk RNA and ATAC-seq time points and conditions for: normoxic (6, 8, and 10 hpf), acute and prolonged paused (6 hpf + 2 hrs hypoxia, 6 hpf + 18 hrs hypoxia, and 6 hpf + 18 hrs KCN (RNA-seq only)), and acute and prolonged reoxygenated (6 hpf + 18 hrs hypoxia+ 1 hr reoxygenation, 6 hpf + 18 hrs hypoxia+ 4 hrs reoxygenation) embryos. PCA plot of bulk RNA-seq samples, see also SSA. Arrows indicate proposed dynamics. **(B)** Volcano plot of differentially expressed genes identified between normoxia (6 and 8 hpf) and hypoxia 18 hrs conditions. Select genes are labeled, see also Data S3. **(C)** KEGG pathways displaying significant enrichment (P-adj<0.05) between normoxia 6 and 8 hpf vs hypoxia 18 hrs. Representative genes were manually selected, see also Data S5. **(D)** Scatterplot of differentially expressed gene log2 fold changes (Wald’s test, P-adj<0.01) in paused conditions comparing 18 hrs hypoxia and 6 hpf normoxia vs 18 hrs KCN and 6 hpf normoxia. Linear least-squares regression line in red. See also Data S3.

We next assessed the consistency of the DEEP program across both time and perturbation conditions. Comparison of DGE signatures from embryos exposed to hypoxia- and KCN-induced pausing revealed a moderate correlation (r=0.61, P-value=8.95e-261) and highlighted both perturbation-specific and shared component genes (Fig. 3D, SSE, Data S6). Comparisons between timepoints revealed that DEG patterns in acute (2 hr) and prolonged (18 hr) hypoxia exposure were highly correlated (r=0.92, P-value<0.0e-308), indicating the continuous elaboration of a coherent expression program (Fig. S5F). We similarly examined the dynamics of pausing exit following reoxygenation. Approximately ∼4 times fewer differentially expressed genes (671 vs 2493) were identified in reoxygenated samples than hypoxic (18 hrs) samples, both relative to normal development controls (Fig. S5G, Data S3). Together, these results confirmed that (1) embryos undergo rapid (1-4 hr) and coordinated changes in both transcriptomic and epigenetic states in response to pausing, and (2) upregulated DEEP transcripts exhibit hallmarks of dormancy, including FoxO signaling and a reduction in anabolic processes. In Data S6 and S3, we report the comprehensive list of DEEP-associated genes, along with associated differential expression statistics.

### Paused embryos require *tentSba* for robust survival and axial patterning

Because paused embryos retained cell type identity at the transcriptional level, we reasoned that some DEEP genes might act post-transcriptionally to stabilize cell states during pausing. To identify candidate regulators, we prioritized DEEP genes encoding post-transcriptional regulatory factors that also displayed strong upregulation during pausing. Among these top candidates was *tent5ba (terminal nucleotidyltransferase 5ba),* which was rapidly upregulated following both hypoxia and KCN-mediated pausing (Fig. S7A) in a HIFla-independent manner (Fig. S6A-B, S7C-D). *tent5ba* also resided within the most significantly differentially accessible ATAC-peak detected following hypoxia (Fig. S7B, Data S4). *tent5ba* encodes a protein sharing ∼67% amino acid identity with human TENT5B (Fig. SSA) and belongs to a conserved family of noncanonical poly(A) polymerases (ncPAPs)(20–23). Unlike canonical nuclear-localized poly(A) polymerases, which extend poly(A) tails co-transcriptionally (*24*), ncPAPs including Tent5 extend poly(A) tails in the cytoplasm (*25–31*). Tent5 proteins have established roles in immunity, secretion, bone formation, gametogenesis, and oncogenesis (*20, 23, 25–29, 32–41*) but have not been extensively studied in the context of embryogenesis, pausing, or hypoxia tolerance - roles that we explore here.

To define the potential roles for *tent5ba* in developmental pausing, we used CRISPR-Cas9 to generate a *tent5ba* mutant allele harboring a 42-bp in-frame deletion near the predicted transferase active site (Fig. SSA). Under standard conditions, both zygotic and maternal zygotic (MZ) homozygous mutants appeared phenotypically normal; these embryos completed embryonic and larval development, and matured into fertile adults (Fig. S8B). By contrast, *MZtent5ba*^-42/-42^ mutants subjected to 18 hours of pausing displayed reduced survival relative to wildtype (Fig. 4A, SSC) and developed axial patterning defects after resuming development.

**Fig. 4.**
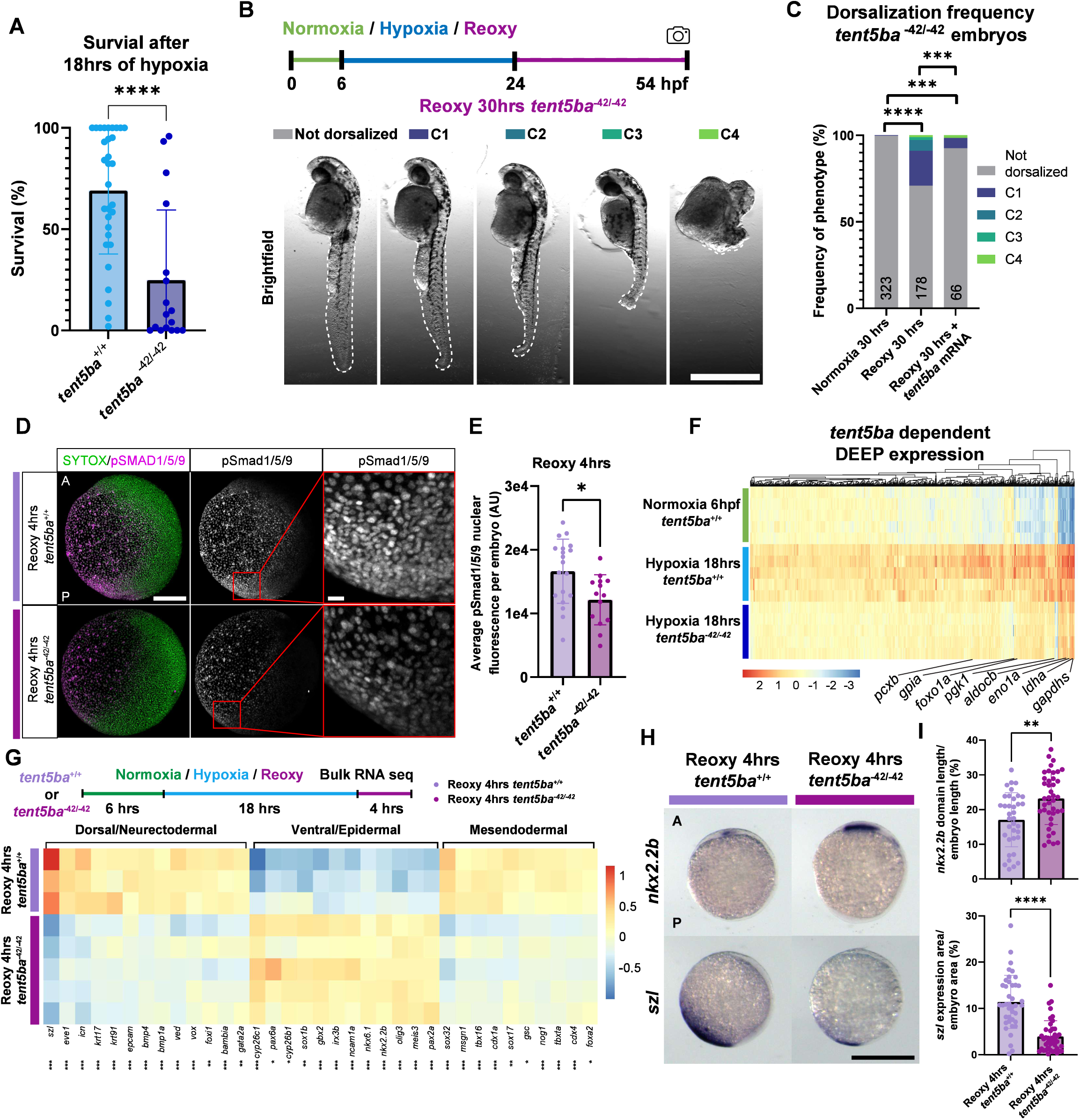
Tent5ba promotes hypoxia tolerance and axis patterning after reoxygenation. **(A)** Survival after 18 hrs of hypoxia, *tent5ba*^+/+^(n=31) and *MZtent5ba*^-42/-42^ (n=l 7) embryos measured after 18 hrs of hypoxia and 24 hrs of normoxia. Dot represents ∼50 embryo groups. Mann-Whitney. See also Fig. SlD and S8C. **(B)** Representative images of dorsalized phenotypic categories observed in *MZtent5ba*^-42/-42^ embryos after reoxy 30 hrs. Scale, 576 µm. **(C)** Proportions of dorsalizations in *MZtent5ba*^-42/-42^ control, reoxy, and mRNA overexpression rescue conditions. Fisher’s exact. **(D, E)** Representative images and quantification (inset) of pSmadl/5/9 labeling in *tent5ba*^+/+^(n=19) and *MZtent5ba*^-42/-42^ (n=14) embryos after reoxy 4 hrs. Mann-Whitney. Scale bars, 154.8 µm or 15.48 µm (zoomed). **(F, G)** Heatmap of column-centered log2-normalized gene counts for (F) genes that were upregulated (P-adj<0.05, log2FC>0.5) in *tent5ba*^+/+^embryos after hypoxia 18 hrs and downregulated (P-adj <0.05, log2FC < −0.5) in hypoxia 18 hrs *MZtent5ba*^-42/-42^ (compared to hypoxia 18 hrs *tent5ba*^+/+^). Glycolysis/gluconeogenesis labeled and (G) axial patterning marker genes in *MZtent5ba*^-42/-42^ and *tent5ba*^+/+^embryos after reoxy 4 hrs, see also Data S7. Asterisks indicate adjusted P-values from DESeq2. (H, I) Representative images and quantifications of whole-mount in situ hybridization of *nkx2.2b* and *szl* expression of *tent5ba*^+/+^(n=36, n=40, respectively) and *MZtent5ba*^-42/-42^ (n=40, n=42, respectively) after reoxy 4 hrs. A: ’Anterior’, P: ’Posterior’. Mann-Whitney. Scale, 410 µm. In all figures, (P or adjusted P-values): * = P≤0.05, ** = P≤0.01, *** = P≤0.001, **** = P≤0.0001 (Methods).

Most prominent among these defects were disruptions to dorsoventral patterning, which included expansion of dorsal neurectoderm and loss of ventral and posterior tail fin structures (*10, 42*) (Fig. 4B-C). We noted that this *tent5ba-dependent* pausing phenotype resembled a more pronounced and penetrant version of the rare (1.3%) dorsalizations observed in wildtype reoxygenated embryos (Fig. SlF-G). Approximately 27% (52/178) of *MZtent5ba*^-42/-42^ embryos displayed aspects of dorsalized morphology, defects that could be rescued by *tent5ba* mRNA overexpression (Fig. 4C, S9A). Consistent with these genetic data, embryos injected with antisense morpholino oligonucleotides targeting either the *tent5ba* start codon (ATG MOs) or a 5’ UTR site (UTR MO) also resulted in dorsalized phenotypes after reoxygenation, which were also rescued by *tent5ba* overexpression (Fig. S9B-D). Notably, the dorsoventral transformations observed after *tent5ba* disruption are consistent with a mid-gastrulation disruption of BMP signaling, which preferentially impacts posterior structures (*43, 44*). This suggested that patterning disruptions occurred after the early gastrulation (shield) stage at which embryos entered the paused state. Consistent with this idea, we observed that post-pausing recovery of pSmadl/5/9 signaling states was impaired in ventral posterior regions ofreoxygenated *MZtent5ba*^-42/-42^ embryos, relative to controls (Fig. 4D-E). Collectively, these data indicate that *tent5ba* is required during and following embryonic pausing for both survival and normal resumption of developmental patterning.

### *tentSba* safeguards transcriptional states that promote hypoxia tolerance and axial patterning

To examine the molecular basis for these patterning and survival defects, we profiled bulk gene expression in *MZtent5ba*^-42/-42^ embryos during both pausing and reoxygenation. To better understand the basis for reduced survival rates, we first examined the impacts of *tent5ba* disruption on DEEP expression. Of the 3,587 transcripts induced under hypoxia in wildtype 6 hpf embryos, 651 (18.1%) were significantly reduced *inMZtent5ba*^-42/-42^ mutants (Fig. 4F, Data S7). This *tent5ba-dependent* subset was emiched for genes within glycolysis/gluconeogenesis pathways, including *ldha, enola, andpgkl* (Fig. 4F, SlOA, Data S5, S7). Expression of *ndrgla* and *ndrgl b,* conserved hypoxia-responsive genes that enable hypoxia tolerance, was also significantly reduced (Fig. S10B-C)(*45*). These transcriptional changes were accompanied by reduced L-lactate levels in paused *MZtent5ba*^-42/-42^ mutants (Fig. S10D), indicating a failure to properly adapt to the hypoxic challenge.

We next examined the expression profiles of early-stage recovering *MZtent5ba*^-42/-42^ embryos using panels of marker genes expressed by and/or required for the specification and patterning of key embryonic lineages (*13–15*). Consistent with subsequent dorsalization, *MZtent5ba*^-42/-42^ mutants displayed coordinated upregulation of dorsal neurectoderm marker genes *(meis3,pax2a, gbx2, nkx2.2b, olig3, ncamla)* and dowmegulation of ventral and epidermal markers *(szl, bmp4, krtl* 7, *ved, epcam, icn)* (Fig. 4G, Data S3). These trends were further validated by whole-mount ISH labeling of *szl* and *nkx2.2b,* whose expression domains were reduced and expanded, respectively, in reoxygenated *MZtent5ba*^-42/-42^ mutants (Fig. 4H-I). In addition, we also noted upregulation of canonical epiblast markers *(pou5j3, klfl* 7, *sox2,* and *myca)* and concomitant dowmegulation of mesoderm *(tbxta, tbxl 6, msgnl)* and endoderm *(sox32, soxl 7,foxa2)* fates, transformations that suggest defects in germ layer induction in addition to those of canonical dorsoventral patterning (*46, 47*) (Fig. 4G, SlOE). Together, these data indicate a broad pleiotropy of developmental defects in paused *MZtent5ba*^-42/-42^ embryos that impact hypoxia tolerance, dorsoventral patterning, and germ layer specification.

### TentSba promotes transcriptome maintenance and extends poly(A) tails of key targets

We next explored the possibility that Tent5ba promotes pausing-induced stabilization of transcriptional programs through non-canonical polyadenylation. Poly(A) tail length (PAL) is a key determinant of mRNA stability as deadenylation promotes decapping and degradation of cytoplasmic transcripts. As transcripts age, PAL serves as a “molecular timer” that tracks degradation dynamics(31, *48–50*); PAL extension, by contrast, could lead to transcript stabilization. To explore this hypothesis, we first examined RNA-seq reads mapping exclusively to introns (pre-mRNAs) or to exons (mature mRNAs) for all genes(51). This analysis revealed that pausing led to an emichment of mature mRNA relative to pre-mRNA for most genes (Fig. SA, Sl lA, Data S8), consistent with global transcriptome stabilization in the paused state. To determine whether these trends correlated with polyadenylation status, we implemented Nano3P-seq(52, *53*), a long read 3’end-cDNA sequencing method to quantify PAL in wildtype and *MZtent5ba*^-42/-42^ mutant embryos (Fig. 5B). Initial control experiments demonstrated that transcript-specific PAL measurements obtained from wildtype 6 hpf zebrafish embryos were highly concordant with previously published zebrafish Nano3P-seq data from the same stage (r = 0.73, P = 5.46e-252; Fig. Sl 1B)(52). We next asked whether and how embryonic pausing affects the global PAL landscape. Hypoxic pausing led to widespread reductions in the mean and variance of PALs across the vast majority (91.7%) ofreliably sampled transcripts (Fig. 5C, S11C). These reductions in PAL were generally not correlated with changes in gene expression, except for a set of 66 transcripts that displayed significant increases in both PAL and transcript abundance. These transcripts included several highly ranked DEEP genes *(hsp70.3, enola)* and *tent5ba* itself, and were also enriched for glycolysis and gluconeogenesis pathway components *(aldoaa, gpia, enol a, pgkl)* (Fig. 5C, Table S5). Global decreases in PALs during pausing - except for select DEEP genes - are indeed consistent with the notion that PALs reflect an increased age of most transcripts in paused embryos. This result, together with the global enrichment of mature, spliced mRNA, suggests that embryonic pausing correlates with long-term maintenance of an early gastrula-stage-like transcriptome.

**Fig. 5.**
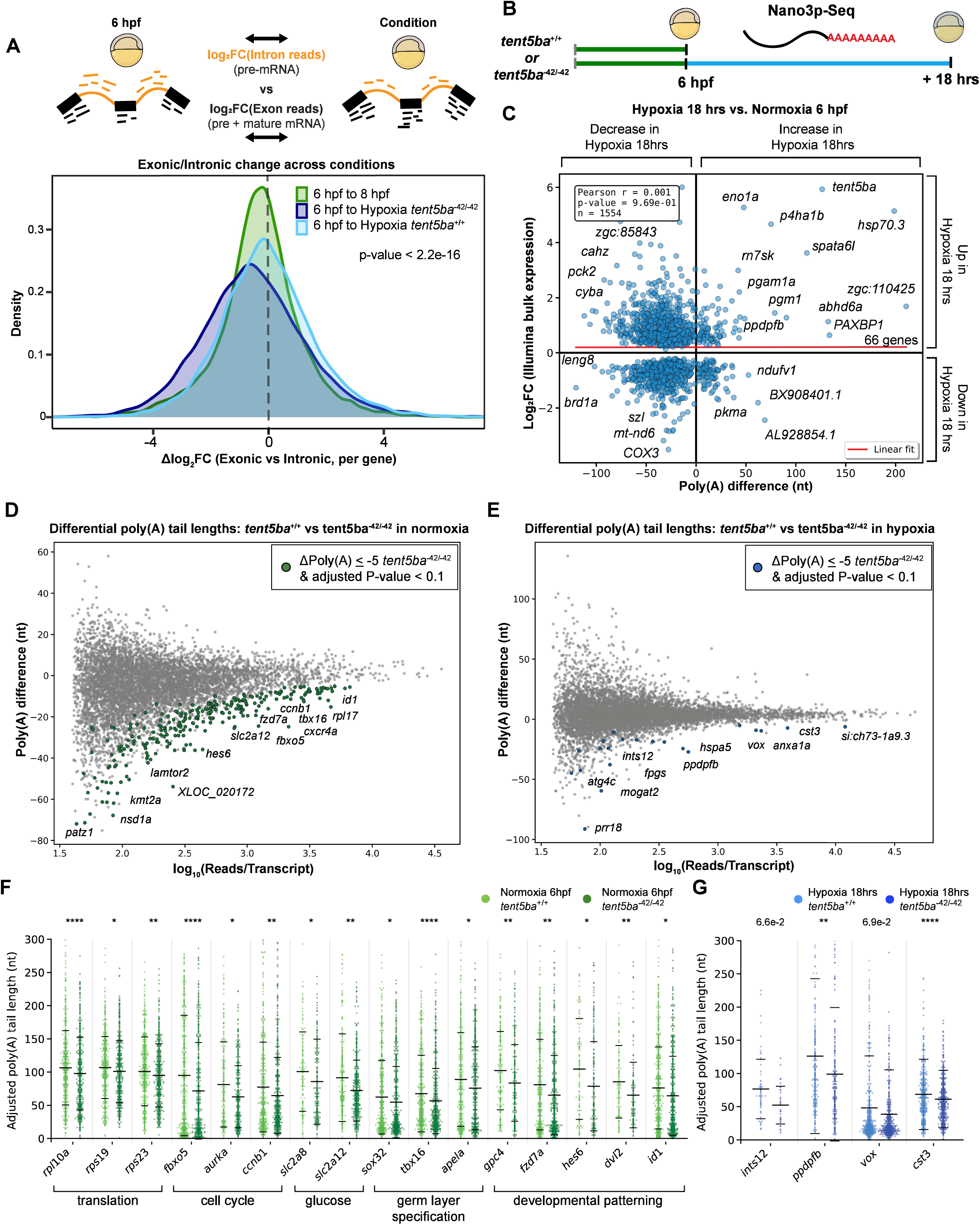
Pausing and *tentSba* post-transcriptional regulation. **(A)** Top: Schematic of bulk RNA-seq Exon-Intron Split Analysis. Bottom: Kernel density estimate of per gene differences between log2FC(Exons across conditions) and log2FC(Introns across conditions). Positive and negative values indicate enrichment towards mature or pre-RNA, respectively. Kruskal-Wallis test, see Data S9. **(B)** Summary ofNano3p-Seq time points and conditions **(C)** Differential polyadenylation (P-adj≤0.05, from Nano3p-seq) vs gene expression (P-adj≤0.05, from Illumina-based bulk RNA-seq). Select genes are labeled, see also Data S3, S8. **(D, E)** From Nano3p-seq, transcripts displaying significantly reduced polyadenylation (P-adj≤0.1, change in nucleotides<-5) by log10 quality-filtered reads. **(F, G)** Beeswarm plots of PAL measurements in control vs *MZtent5ba*^-42/-42^ mutants in standard conditions (green) and hypoxia (blue). Mean and standard deviations are indicated by black lines. PAL data are plotted for range 0-300 nt. See also Data S8. In all figures, (P or adjusted P-values): * = P≤0.05, ** = P≤0.01, *** = P≤0.001, **** = P≤0.0001 (Methods).

We reasoned that a plausible function for *tent5ba* could be to promote transcript stabilization during the paused state, perhaps via post-transcriptional PAL regulation. Indeed, paused *MZtent5ba*^-42/-42^ mutants displayed a reduced proportion of mature mRNA relative to pre-mRNA, suggesting global transcriptome destabilization), lingering transcription, or a combination of the two (Fig. 5A, Sl lD). Consistent with *tent5ba* acting as an ncPAP, *MZtent5ba*^-42/-42^ mutants developing under standard conditions displayed reduced PALs (Fig. 5D, S1lE), with a modest positive correlation between changes in PAL and differential expression (r=0.32, P=0.02, n=51) (Fig. S11F). Transcripts that exhibited decreased PAL in *MZtent5ba*^-42/-42^ mutants were enriched for ribosome, spindle, and nuclear components, as well as translation *(rpll Oa, rps] 9, rps23),* cell cycle *(jbxo5, aurka, ccnbl),* glucose metabolism *(slc2a8, slc2al 2),* germ layer specification *(sox32, tbxl 6, apela),* and developmental patterning *(gpc4,fzd7a, hes6, dvl2, idl)* (Fig. 5D-E, Data S5)- several of the same processes that manifested later as post-pausing developmental defects. In paused embryos, additional *tent5ba-*dependent PAL signatures were also identified for *cst3,* the homologue of a recently reported TENT5A/C target(20, *28*), as well as *intsl 2* and *ppdpjb,* components of the integrator complex(54) and mTOR pathways(55), respectively, and *vox,* a homeobox transcription factor known to promote ventrolateral fates(56–59) (Fig. 5E-F). These data indicate that *tent5ba* indeed functions as an ncPAP and reveal that several of its downstream transcriptional targets promote developmental processes that are disrupted in paused and/or recovering *MZtent5ba*^-42/-42^ mutants. We conclude that Tent5ba promotes successful developmental pausing through poly(A)-tail extension and transcript stabilization and/or maintenance, buffering a latent vulnerability imposed by reliance on long-lived transcripts during pausing.

## Conclusion

Here, we describe embryonic pausing as an adaptive resilience strategy that preserves developmental fidelity under extreme environmental stress. Upon pausing, zebrafish embryos undergo a striking decoupling of signaling state and transcriptional identity: canonical morphogen gradients collapse yet core transcriptional features of embryonic identity are largely maintained. Successful pausing relies on the expression of a protective gene expression program (DEEP), which includes *tent5ba,* a conserved non-canonical poly(A) polymerase, as a critical factor. Tent5ba promotes DEEP expression, hypoxia tolerance, transcriptome maintenance, and faithful developmental patterning post-pausing. We propose that poly(A)-tail based post-transcriptional regulation, mediated in part by Tent5ba, constitutes an adaptive buffering mechanism that safeguards embryonic transcriptomes through periods of suspended embryogenesis.

## Acknowledgments

We thank Y. Su and N. Suren for zebrafish husbandry; P. Bhat, K. Chen, C. Chou-Freed, I. Roy, N. Aponte Santiago, H. Smith, and **J.** Zussman, for helpful discussions; R. Boileau for ATAC-sequencing advice. J. Reiter, I. Jain, and P. Singh for scientific guidance, H. Smith and A. Cesiulis for in situ plasmid help, and R. Sahasrabudhe for Nanopore-sequencing advice.

## Author contributions

Conceptualization: CPC, DEW

Methodology: CPC, HG, DEW Investigation: CPC, SF Visualization: CPC

Funding acquisition: CPC, DEW Project administration: CPC, DEW Supervision: DEW

Writing - original draft: CPC, DEW

Writing - review & editing: CPC, HG, SF, DEW

## Data, code, and materials availability

Raw and processed data will be made available through the NCBI Gene Expression Omnibus (GEO) and Sequence Read Archive (SRA) repositories, or by request.

## Supplementary Materials

Materials and Methods

Figs. Sl to Sll

Data Sl to Sl 1

References (60–94)

## Materials and Methods

### Zebrafish Husbandry

Unless otherwise stated, all Zebrafish *(Dania rerio)* experiments conducted were on the background of AB strains from ZIRC. All practices were approved by the University of California, San Francisco’s Institutional Animal Care and Use Program and monitored regularly by the Laboratory Animal Resource Center. Animals were housed in the Wagner lab’s fish facility at 28.5°C on a 14 hr - 10 hr light-dark cycle. Embryos were acquired via timed matings and kept at 28.5°C in 0.3x Danieau buffer (17.4 mM NaCl, 0.21 mM KCl, 0.12 mM MgSO4, 0.18 mM Ca(NO3)2, 1.5 mM HEPES, pH 7.6). Embryos were staged by wall clock time, assisted by classic morphological criteria(9). When necessary, embryos were dechorionated by incubating them in 1 mg/mL Pronase (Sigma-Aldrich, 53-702-500KU) for 5 minutes, followed by three washes with 0.3x Danieau buffer. All experiments were conducted before zebrafish undergo sex determination (∼25 days post-fertilization)(60).

Zebrafish lines and their generation with CRISPR

In this study, we generated a stable mutant *tent5ba* zebrafish line *(MZtent5ba*^-42/-42^ using the Alt-R CRISPR-Cas9 System (IDT). Two custom crRNAs (AAGTTTCCACGTCCGTGAAC, TTTACCTGAACAATCTGTCG) were designed on IDT’s Alt-R CRISPR-Cas9 System webpage; guides were selected based on their On-Target score, Off-target score, and their proximity to the TSS. crRNAs and tracrRNA (IDT, 1072532) were resuspended in Nuclease-free IDTE buffer (IDT, 11-01-02-02) at 100 µM each, aliquoted, and stored at −80°C until use. crRNA, tracrRNA, Nuclease-Free Duplex Buffer (IDT, 11-01-03-01) were combined to a final gRNA concentration of 6 µMand heated to 95°C for 5 minutes. Alt-R S.p. HiFi Cas9 Nuclease V3 (IDT, 1081061) was diluted in Cas9 working buffer (20 mM HEPES; 150 mM KCl, pH 7.5) to a final concentration of 1.0 µg/µL. gRNA and Cas9 solutions were mixed together at a **1:**1 ratio, incubated for 10 minutes at 37°C, and allowed to cool to room temperature. Wildtype AB embryos at the one-cell stage were collected and injected with ∼0.5 nL of the ribonucleoprotein mixture with 0.03% phenol red (Sigma, P0290). Genomic DNA was extracted using the HotShot DNA isolation as previously described(61). Editing efficiency in F0 injected embryos was assessed using the T7 endonuclease I (T7El) mismatch cleavage assay (IDT, 1075932) following the manufacturer’s guidelines, on a 211 bp PCR product generated using the primers specified below and Q5 High-Fidelity 2x Master Mix (NEB, M0492L).

Once F0 injected embryos reached sexual maturity, they were outcrossed to nonsibling AB zebrafish to obtain non-mosaic Fis. These Fis were fin clipped using our IACUC’s standard procedure https://iacuc.ucsf.edu/sites/g/files/tkssral6261/files/wysiwyg/STD%20PROCEDURE%20-%20Aquatic%20-%20Fin%20Clipping%20of>/o20Zebrafish.pdf, fish with similar T7 patterns were in-crossed together, and their progeny were genotyped using Sanger sequencing with the same primers (TTGTGTGTTGTCTTGGGAGCA, GTCATGAAGTAGCATTAATAGCCTCAAAG) to assess homozygosity. The exact indel produced in the *tent5ba-^42^* allele can be found in Fig. S8A. Double Hifla mutants *(MZhiflaa* ^bns89/bns89^_;_ *MZhiflab* ^bns90/bns90^, referred to in Main Text as *hifl aa^-/-^ hifl ab^_/_^)* were generously shared with us from Kathleen Gilmour’s lab at the University of Ottawa and were originally generated in the lab of Didier Stainier(*62*)*. hifl aa* and *hifl ab* alleles were amplified by PCR and Sanger sequenced *(hifl aa* primers: CTCCCCTGATTCACCAAAGCATCGG, AATGGGGGCTCGTCTGGGATTTGAA; *hifl ab* PCR primers: AATTCAATGTTGCGGCAAACATGCC, CCGCAGGCTTTTGTGCTGACACTAA) at Azenta Life Sciences. All mutant alleles assayed in this study are maternal zygotic (MZ) mutants. The only exception is for experiments when heterozygotes were used, in which case, *tent5ba^-42/+^* adults were crossed together, and their embryos were used.

### Hypoxia and KCN conditions

For all hypoxia experiments, hypoxia chambers were created from OrnniTop 50 mL Sample Tubes (Avantor, 75840-772). Sample tubes containing 5 mL of 0.3x Danieau buffer were flushed with N2 gas for 15 minutes, three times: 24 hours, 6 hours, and 15 minutes prior to embryo addition, and once after for 15 seconds, each at a flow rate of∼ 12 bubbles per second.

Dechorionated embryos at the shield stage survived extremely poorly due to mechanical shearing from N2 flushing. We found that pausing was the least variable when 50-80 embryos were used per tube; fewer embryos yielded inconsistent results. Sample tubes were stored in incubators set to 28.5°C throughout the process, except during active nitrogen flushing, which occurred at room temperature. Each sample tube was equipped with an oxygen sensor spot (PresSens Precision Sensing, SP-PSt3-NAU-D5-YOP Planar Oxygen-Sensitive Spot, 200000023) that remained fully submerged throughout the experiment. Dissolved oxygen was measured optically using a Fibox 4 Oxygen Meter (PreSens Precision Sensing, 200001478). Oxygen concentration measured immediately following flushing consistently read as ∼0 mg/L before embryo addition. Tubes were measured at the end of hypoxic exposure; tubes that exceeded 1.0 mg of 02 per liter at the end of the hypoxic period were excluded from studies unless explicitly desired (ie, Fig. S1D and S8C).

For all potassium cyanide (KCN) experiments, chorionated embryos were exposed to 1 mM KCN (Sigma-Aldrich, SIAL-207810) dissolved in 0.3x Danieau buffer. KCN is volatile, which means that 1) since it is toxic, it was handled and disposed of with caution and under a hood, 2) vessels with KCN and embryos were sealed with parafilm to keep the effective concentration of KCN consistent throughout the experiment, and 3) conditions containing KCN were kept separate from other experiments because simple proximity was sufficient to pollute non-KCN conditions. After embryos were removed from KCN conditions, they were washed three times for 5 minutes with 0.3x Danieau buffer, very gently shaking at room temperature. For all pausing (KCN and hypoxia) experiments, dead or dying embryos were excluded from further analysis. EC50 values for the hypoxia tolerance panels, were determined in GraphPad Prism (10.4.2) where a nonlinear regression (EC50 shift, xis concentration) was used to fit a curve to dose-response data. Specifically, a least-squares regression was used, and the top and bottom values were constrained to 100 and 0, respectively, to reflect the bounds of the applicable survival data in percentages, see Data S10.

For crowding experiments, embryos were spawned as described above. Six hours after fertilization, 50 embryos were placed into either 1) a 10 cm petri dish with ∼30 mL of 0.3x Danieau buffer or 2) a 1.5 mL Eppendorftube with **1** mL of 0.3x Danieau buffer. Embryos were incubated in either condition for 18 hrs and then imaged using bright-field microscopy as described below.

### Phenotypic Evaluations

Live embryos were examined at various time points using brightfield microscopy on a Zeiss Stemi 305. Images were captured using a Leica M205 FCA stereoscope after embryos were fixed in 4% paraformaldehyde in PBSTx (0.1% Triton X-100, in PBS) with SYTOX green **(1**:20,000) overnight at 4°C rocking. To classify embryos into dorsoventral phenotypic bins, we relied on existing well-established criteria(*10, 42*). In combination with these previous criteria, we specifically focused on the trunk and tail of the embryo. We defined class 1 (Cl) as having a reduced ventral tail fin, class 2 (C2) as a complete reduction in part of the ventral tail fin, class 3 (C3) as having at least a partial reduction in the tail, and class 4 (C4) as a near complete absence of the tail. When discerning between the extent of dorsalization across conditions, a Fisher’s exact test was performed, raw data and statistics are available in Data S10.

To quantify epiboly progression, chorionated embryos were fixed in 4% paraformaldehyde in PBSTx (0.1% Triton X-100, in PBS) with SYTOX green **(1**:20,000) overnight at 4°C rocking. Embryos were mounted in molds, and both brightfield and fluorescent images were captured using a Leica M205 FCA stereoscope. Exported images were blindly quantified using Fiji ImageJ (version 1.45p), where the length of the epiboly extent was measured from the tip of the animal axis to the furthest cells reach along the vegetal pole, excluding yolk syncytial cells. This length was divided by the distance from the tip of the animal pole to the end of the vegetal yolk pole, and the result was converted to a percentage. Within each batch ofreplicates, a Kruskal-Wallis test was performed, raw data and statistics are available in Data S10.

### Immunofluorescence and fluorescence microscopy

For fixation and storage, embryos were incubated overnight in 4% paraformaldehyde, 0.1% Triton X-100, in lx PBS, with the chorions intact, rocking at 4°C. Embryos were moved into PBSTx, manually dechorionated using forceps, and washed three times in PBSTx at room temperature. For storage, embryos were transferred into 100% methanol in a stepwise manner and stored at −20°C. To start staining, embryos were rehydrated in a stepwise manner back to lx PBSTx.

Imrnunofluorescence was performed similarly to as previously described(63, *64*). In brief, samples were blocked in NCS-PBST (10% fetal bovine serum, 1% DMSO, 0.1% Triton X-100 in PBS) for 2 hours, incubated overnight in primary antibodies in NCS-PBSTx rocking at 4°C, and then incubated overnight in secondary antibodies (Invitrogen, Al 1001 and A21245, **1**:500) in NCS-PBSTx rocking at 4°C with either SYTOX green (Invitrogen, S7020, **1**:20,000) or DAPI (Invitrogen, D21490,1:10,000). To aid imaging efficiency and accuracy, embryos were flat-mounted after staining for pSmadl/5/9 (Cell Signaling, 13820. 1:200), β-catenin (Sigma Aldrich, C7207, **1**:1000), and pErkl/2 (Cell Signaling, 4370, **1**:500). First, the embryos were deyolked as previously described(63), and then cut four times inwardly (from margin to animal) at 90° intervals, approximately two-thirds of the way towards the animal cap, using 2 mm Vannas spring scissors (Fine Science Tools, 15000-03). Our flat-mounting procedure resembles that of flat-mounted retinas(65). For the pSMAD2/3 (Cell Signaling, 8828, 1:800), after rehydration, embryos at 4 hpf were deyolked and incubated in precooled acetone at −20°C for 20 minutes. To stain embryos injected with mRNA containing the ALFA tag (see below for more information), embryos were incubated overnight with an ALFA-647-conjugated antibody (NanoTag Biotechnologies, N1502-AF647-L, 1:500) as above, without a secondary antibody incubation. Before imaging, all embryos were mounted into ProLong Gold Antifade Mountant (Invitrogen, P36930). Immunostained embryos were imaged on a Leica Stellaris 5 laser scanning confocal microscope using either an HC FLUOTAR L 25x/0.95 W VISIR or an HC PL APO 10x/0.40 CS2 objective (Leica).

### Morphogen gradient quantification

For the morphogen gradients quantified in 4 and 6 hpf staged embryos, images were acquired across the entire embryo. Images were imported into Imaris (Oxford Instruments, version 10.2.0), and a region of interest was segmented by generating a ∼250 µm wide volume spanning the margin-to-margin axis. A new reference plane was used to redefine the Cartesian coordinates of the volume, with one side of the margin at (0,0). To identify nuclei, the spots tool was applied to the SYTOX or DAPI channel, assuming a nucleus diameter of 7 µm, with no background subtraction and using semi-consistent quality filters. To specifically quantify the nuclear beta-catenin signal, a surface was generated from the SYTOX or DAPI channel that excluded the cytoplasmic staining. The intensity mean of every spot was exported into R (version 4.3.1), along the margin-to-margin axis nuclei were binned into deciles, and their fluorescence intensity was averaged. Ggplot2’s (version 3.5.2) geom_smooth function (method= “loess“, Local Polynomial Regression Fitting) was used to plot the average fluorescence intensities aggregated across replicates. For each decile, a Wilcoxon T test was performed, and the significance was displayed as asterisks above. Because the differences between normoxic and hypoxic samples were so large, it was not possible to blind investigators during quantification.

### Raw data and statistics are available in Data S10

For the pSmadl/5/9 quantification during reoxygenation (4 hrs), images were acquired encompassing one lateral half of the embryo. Images were imported into Imaris software, where a region of interest (350 px by 350 px by Max Z) was placed within the ventral tail bud, and the region was segmented by creating a new volume. Nuclei were identified by using the spots tool on the SYTOX Green channel, assuming a diameter of 6 µm with background subtraction and using semi-consistent quality filters. The pSmadl/5/9 fluorescence intensity sum for each nucleus was exported and averaged per embryo. A Mann-Whitney test was performed between conditions. When semi-subjective portions of this analysis were conducted, such as the placement of the region of interest, the sample names were hidden from investigators. Raw data and statistics are available in Data S10.

### Single-cell dissociations, library preparation, and sequencing

Single-cell dissociations were adapted from previously reported methods(66). Dissociations were conducted in precoated tubes prepared by adding 1% Fraction V BSA (Gibco, 501215315) to 1.5 mL Protein LoBind Tubes (Eppendorf, 13-698-794) for 30 minutes at room temperature. The tubes were then decanted and left to evaporate in a biosafety cabinet for an additional 30 minutes at room temperature. Approximately 50 embryos were added to each tube. Then, 150 µL of ice-cold 1% Fraction V BSA in PBS was added, and embryos were dissociated by pipetting 10 times using a P200 pipette set to 150 µL. Cell suspensions were then fixed using Parse Biosciences Evercode V2 Fixation Kits (ECF2101) and brought gently to - 80°C in a CoolCell LX cell freezing container (Coming, 432138). Before freezing, suspension concentrations were quantified using a hemocytometer.

Single-cell barcoding and library preparation were conducted using the Evercode WT v2 Barcoding Assay (Parse Biosciences, ECW020301). We followed the manufacturer’s instructions for the majority of the protocol. For the amplification of barcoded cDNA we used 6 PCR cycles for the number of 2nd cycles (X), and for the sublibrary index amplification, we used 9 PCR cycles (X). Sublibraries 1 and 2 of 8 were first sequenced on an Illumina NovaSeq X (2 Lanes, 1.5 billion clusters, il:8, i2:8, Rl:136, R2:86, PhiX:5%) to first assess the quality of cells. The remaining 6 of 8 sublibraries were sequenced on another Illumina NovaSeqX (5 Lanes, 6.3 billion clusters, il :8, i2:8, Rl :100, R2:100, PhiX:15%), also at the Chan Zuckerberg Biohub San Francisco.

Raw FASTQs were sent through the Parse Biosciences Pipeline bioinformatics program split-pipe (version 1.0.3p) to generate an unfiltered cell-gene count matrix. Reads were aligned to an indexed reference genome made using split-pipe from the genome assembly GRCzl 1 and the Lawson Lab transcriptome annotation V4.3.2.gtf(67). This was imported into scanpy (version 1.11.5), cells were only retained if they exceeded 4500-5000 reads per barcode and had a low doublet score (scrublet, version 0.2.3, less than *0.1)*(*68, 69*). Cells were removed if more than 50% of their reads arose from mitochondrial genes; however, fewer than 1-2% of reads mapped to mitochondrial or ribosomal genes across all cells. Quality metrics for filtered cells are provided in Data S1.

### Single-cell cell type annotations

In summary, cells were assigned cell type annotations using a CellTypist (version 1.6.3) classifier trained on normoxic wild-type cells generated in this study(70). To generate the training dataset, only normoxic cells (6, 8, and 10 hpf) were used. Raw counts per cell were log-transformed (log(l +x)) and normalized by read depth (TPM). The scanpy function highly_variable_genes was used to identify the top 2000 variable genes in each replicate before merging them. The number of non-trivial principal components was selected by juxtaposing the eigenvalue distribution generated from randomized data, as previously described (7l)(https://github.com/wagnerde/scTools-py). Leiden clustering was performed at varying resolutions, then rank_genes_groups using Wilcoxon rank-sum and Benjaminin-hochberg p-value correction. Additionally, we performed a cell annotation label transfer from a comprehensive zebrafish developmental atlas (subset to include cell types between 6-10 hpf) called ZMAP using Symphony(*15, 72*). Together, we used these annotations to inform our Leiden resolution (0.75) and cluster nomenclature, which we found highly resembled other single-cell datasets and matched classic cell-type marker gene expression patterns . We used this subset of the dataset and its annotations to train a CellTypist model. We applied this model to our entire single-cell dataset to uniformly annotate cells across all conditions. To visualize samples from all conditions together, quality-filtered and annotated datasets were aggregated, processed as described above, and harmonized (version 0.010) by condition(73). Neighbors (10) were calculated using harmonized PCA space and embedded with Uniform manifold approximation and projection (UMAP)(74).

### Single-cell Euclidean distance measurements

Quality-filtered and annotated datasets were subset to include only samples and cell types being directly compared. Principal component analysis and neighborhood connectivity were recalculated as described above. The pairwise Euclidean distance across principal component space was measured using PertPy’s (version 1.0.3) distance.pairwise function (75) across cells annotated by their cell type, condition, and replicate. These distances were binned into two categories: 1) within normoxic cells, the average distance between a specified cell type and all other normoxic cell types (excluding its own cell type), and 2) the distance between a specified cell type in hypoxia and normoxia. These distance measurements were repeated for each replicate and cell type, and the results were compared using a two-sided Mann-Whitney U-test, these data are available in Data S10.

### Single-cell Differential Gene Expression Analysis

Differential gene expression analysis was conducted by psuedobulking cell types and replicates. To do this, we implemented a Python pseudobulk version of DESeq2, called PyDESeq2(75).

Differentially expressed genes (padj < 0.05, llog2foldchangel > 0.5) identified by this method were compared to the global hypoxia CNMF GEP, 2000 of the top-ranked CNMF GEP5 genes (see below). A table of genes differentially expressed by their cell type can be found in Data S2.

### Single-cell Gene Expression Programs

Gene expression programs (GEPs) were identified using consensus non-negative matrix factorization (cNMF) (version 1.7.0)(16). The number of components was determined empirically by calculating stability and error across multiple k selections, iteratively over increasingly selective windows. Eventually, a k of 40 was identified as being the most stable with the least error. The k-selection was conducted utilizing the top 2000 most variable genes. We ran cNMF with 40 components to generate a cell usage matrix and top z-scored genes for each GEP. We discerned between ’identity’ and ’activity’ GEPs by how specific the GEP usage score was to either previously annotated cell types or across our experimental conditions. A table of the genes contributing to each cNMF gene expression program is available in Data S1.

### Bulk RNA sequencing

We performed bulk RNA sequencing using two separate technologies 1) 3’ Tag-Seq prepared and sequenced by the UC Davis Genome Center at the DNA Technologies and Expression Analysis Core (referred to as “TagSeqA” and “TagSeqB“) and 2) mRNA sequencing (referred to as “mRNASeq“) prepared and sequenced by Novogene. The RNA extraction protocols for both technologies began similarly, with RNA extracted using TRizol Reagent (Invitrogen, 15596026), following the manufacturer’s protocol for the most part. Embryos at the specified stage/condition were dechorionated, lysed in TRizol with vigorous pipetting using a P200 pipette, and stored at −80°C for future use. Lysates were phase separated by the addition of chloroform. For 3’ Tag seq, the aqueous layer was added to an RNA Clean & Concentrator-5 spin column (Zymo Research, R1013) and subjected to a DNase I treatment on the column. For mRNA-seq, RNA was precipitated from the aqueous phase by adding isopropanol, sodium acetate, and GlycoBlue Coprecipitant (Invitrogen, AM9516). It was then washed with ethanol and reconstituted in RNase-free water. Both samples were assessed on an Agilent TapeStation 4200 using an RNA ScreenTape (Agilent, 5067-5576). Samples for 3’ Tag-seq were checked for contaminants on a NanoDrop One (Thermo Scientific). Library preparations for 3’ Tag Seq were conducted with UMI tags and single-end sequenced on either a NextSeq500 or Element Bio Aviti sequencer at the UC Davis Genome Center. mRNA-seq samples were submitted to Novogene, where poly(A) capture enrichment, cDNA reverse transcription, and 150 bp paired-end sequencing were performed on the Illumina X plus platform. Sample information is available in Data S3.

All samples were processed using the Nexflow (version 23.04.2, (76)) nf-core maseq (version 3.12.0, (77)) pipeline. Described briefly, reads were aligned and quantified with Star (version 2.7.9a, (78)) and Salmon (79) using GRCzl 1/danRerl 1 version of the zebrafish genome with the Lawson Lab’s transcriptome annotation (v4.3.2)(67). Gene count matrices were imported into DESeq2 (version 1.40.2, (80)) for differential gene expression analysis. Principal component analysis was performed and plotted with the plotPCA function ofDESeq2. The results of the PCA analysis were used to identify the best comparisons for differential gene expression tests, thereby minimizing staging and technical batch effects. Volcano plots were made with EnhancedVolcano (version 1.18.0, (*81*)), log2FC values presented in volcano plots are shrunken log fold changes. Gene set enrichment analysis and Overrepresentation analysis were conducted using clusterProfiler (version 4.8.3, (*82*)). Differential gene expression tables can be found in Data S3.

Exon lntron Splice Analysis (EISA) was conducted as previously described (*51*). In brief, reads were aligned to the GRCzl 1/danRerl 1 version of the zebrafish genome with the Lawson Lab’s transcriptome annotation (v4.3.2) using Rhisat2 (splicedAlignment = TRUE, version 1.16.0). Both exonic and intronic reads were quantified via qCount from QuasR (1.40.1), intronic counts were obtained by the difference between gene bodies and exonic counts. Exon and intron counts were normalized to the mean library size separately and logarithmically transformed with a pseudocount. Genes with low counts were filtered out. Differential testing was conducted to compare exonic and intronic changes across conditions using eisaR (1.12.0), which utilizes edgeR’s (3.42.4) model. Exon/intron ratios for each library and differential testing results can be found in Data S9, S10.

### Bulk ATAC sequencing

Embryos were collected at the defined stages/conditions, dechorionated, and dissociated as described in the “Single-cell dissociations, library preparation, and sequencing“. After one round of washing and centrifugation in 1% Fraction V BSA in PBS, cell suspensions were resuspended in 50 uL of CryoStor cell cryopreservation media (Sigma-Aldrich, C2874). 5 uL of the suspension was used to approximate cell number via a hemocytometer. The remainder of the suspension was brought to −80 °C gently in a CoolCell LX cell freezing container (Corning, 432138). Tagmentation and library preparation were performed on ∼65,000 high-quality cells per sample using the Active Motif ATAC-Seq kit (53150) according to the manufacturer’s protocol. Libraries were quantified on a Qubit 2.0 Fluorometer (lnvitrogen, Q32866) using a dsDNA Quantification Assay Kit (lnvitrogen, Q32851) and assessed on an Agilent Tapestation 4200 using a Dl000 Screen Tape (Agilent, 5067-5582). Libraries were sequenced using an Illumina NextSeq 2000 P3 **(1.**1 billion clusters) platform (il :8, i2:8, Rl :61, R2:61, 1% PhiX) at the Chan Zuckerberg Biohub San Francisco.

FASTQs were processed using the Nextflow (version 23.04.2) on the nf-core atacseq workflow (version 2.0). Briefly, Trim Galore (version 0.6.7) was used to trim adapters (*83*). Alignment was conducted using BWA (version 0.7**.1**7-r1188, (*84*)) to the zebrafish reference genome GRCzl 1/danRerl 1 version of the zebrafish genome with the Lawson Lab’s transcriptome annotation (v4.3.2)(67). Narrow peaks were called with MACS2 (version 2.2.7.1, (*85*)). We performed differential accessibility analysis using Diffbind (version 3.10.1, (*86*)) following previously established protocols (*87*). Peaks were assigned to their nearest gene (-2000 to +500 bp from a TSS) using ChIPseeker (version 1.36.0, (*88*)). Reads were normalized and quantified per accessible region and imported into DESeq2 (version 1.28.0) to generate the PCA plots. Differential accessibility results are available in Data S4.

### Microinjections of one-cell stage zebrafish embryos

Embryos were collected after a timed mating and injected at the one-cell stage. Translation blocking morpholinos were designed and produced by Gene Tools LLC. Sequences are as follows: Standard control (STD MO) – human beta globin intron (CCTCTTACCTCAGTTACAATTTATA), *tent5ba* start codon (ATG MO) (CTGTCGCATCACCGGAGGACATG), *tent5ba* 5’UTR (TTCCAGCAGATGAACTTCCTACGCA). Morpholinos were reconstituted in 0.3x Danieau buffer at 4 mM, warmed to 65 °C for 10 minutes, aliquoted, and stored at −20 °C as previously described (*89*). Injections solutions consisted of 0.1 M KCl, 10% phenol red injection dye, and morpholino. Mixtures were preheated to 65 °C for 10 minutes prior to injection and allowed to cool. Embryos were injected with 0.5 nL of a solution containing 5 ng of morpholino.

To make *tent5ba-ALFA* tagged mRNA, we designed and ordered a gblock from IDT, the sequence is as follows: cctaggcctatttaggtgacactatagaagagtactaatacgactcactatagggagaagctacttgttctttttgcaggatccgccaccatgtc ctccggtgatgcgacagagcagtgtcggcggttttgtgtgttgtcttgggagcaggtgcaacgtttggactctattctcggagagacggttcc agttcacggacgtggaaacttcccgacgctttcggtgcagccgcgacagattgttcaggtggttcgagcacgtttggaagagcgaggaatc acggttcgagatgttaggctgaacggttcagctgccagtcacgttctgcaccaggacacgggcctcggatacaaagacttggacctgattttt ggtgtttctctgtcagatgaccaggctttccgtgtggtgaaggacgtggttctcgacagcctgctggacttcctacctgaaggcgtcagcaaa gagcgaatctcagcactgactctgaaagaggcttacgtgcagaaactggtgaaggtgtgcaacgacaccgaccgctggagcctcatctcg ctttccaataacacaggcaagaacgtcgagctcaagttcgtggactctctaagacggcagtttgagttcagcgtggactccttccagattgtttt ggactctcttcttctgttcgaccgttgttcggagacgcccatgtcggagtgtttccaccctacagttttgggcgagagtatgtatgggaacttcg aggaagctcttggccatctgtgcaacaaggccatcgctacacgaaaccctgaagaaattcgcggaggcggcctcttaaaatactgccacct tttagtgcgaggattccgtcccacttccgaatcggagatgaaatcgctccagcggtacatgtgctctcggttcttcattgactttccagacatcg gggagcagcagcgaaaactggaggcgtacctacagaatcactttgccggaatggagcacaaacggtatgactgtttggtcacgttgtacca cgtggtcaatgagagtactgtatgcttaatgggacacgagcgacggcagactttaaaccttatctcaatgctggcgctaagggtgctcgccg aacagaacgccatccctacagttaccaacgtcacatgttattaccagccggcaccgtacgtgcgagacatcaacttcagtaactactacattg ctcacgtgcagccgctggttcacacttgcagtagttcctatccgacatggctgccctgtaacCCATCACGTCTGGAGGAAG AGCTGCGTCGTCGCCTGACCGAACCTtgatctgcagtaagaattcatacgtatccggaaccggtgatccagacatga taagatacattgatgagtttggacaaaccacaactagaatgcagtgaaaaaaatgctttatttgtgaaatttgtgatgctattgctttatttgtaacc attataagctgcaataaacaagttaacaacaacaattgcattcattttatgtttcaggttcagggggaggtgtgggaggttttttgatatcccggg tttaaacgcatgcagatctaatactcaagtacaattttaatggagtacttttttacttttactcaagtaagattctagccagatacttttacttttaattg agtaaaattttccctaagtacttgtactttcacttgagtaaaatttttgagtactttttacacctctgtcaagaaccatatgccggtacccaattcgcc ctataggacgtcgtattacgcgcgctcactggccgtcgttttacaacgtcgtgactgggaaaacc. In brief, we placed the coding DNA sequence of the *tent5ba* gene (without introns or a stop codon) after an SP6 and T7 promoter, and a KOZAK sequence, and before an ALFA sequence, SV40 poly(A) tail, and an M13 forward sequence. We based this design on other expression vectors (*90*). We received the zebrafish codon optimized sequence of ALFA (CCATCACGTCTGGAGGAAGAGCTGCGTCGTCGCCTGACCGAACCT) from a generous correspondence with Curtis Boswell(91, *92*). We amplified our gBlock via PCR using Sp6 upstream (ATTTAGGTGACACTATAG) and M13 Forward (GTAAAACGACGGCCAGT) primers and cleaned up the product with an SPRI bead clean at 0.6x (Beckman Coulter, A63881). mRNA was made using the mMESSAGE mMACHINE Sp6 Transcription Kit (Invitrogen, AM1340) following the manufacturer’s recommendations. In vitro transcribed mRNA was assessed on an Agilent TapeStation 4200 using an RNA ScreenTape (Agilent, 5067-5576) and was found to be the expected size. mRNA was aliquoted and stored at −80°C for future use. Embryos were injected with 25 pg of mRNA as described above (0.1 M KCl, 10% phenol red injection dye) in a volume of 0.5 nL. For experiments in which embryos were injected with both MO and mRNA, the mRNA was added to the injection mixture after the solution had been warmed to 65 °C. To assess mRNA translation efficiency, injected embryos were stained via the immunofluorescence protocol listed above.

### In situ hybridization

RNA probes for in situ hybridization were generated using gene-specific sequences that were cloned into vectors (pCRII Blunt-TOPO or pGEM-T Easy Vector). Gene-specific sequences were amplified by PCR. RNA probes were made via in vitro transcription off of gene-specific linearized templates using a T3 RNA polymerase (Invitrogen, AM1348) and DIG labeling mix (Roche, 11277073 910). DIG-labeled RNA probes were purified by ethanol precipitation and resuspended in formamide. The size of RNA probes was assayed by running the denatured probe on a 1% TBE gel. Primers and/or accession numbers used to amplify gene specific seuqneces are as follows *szl* (NM_181663.1, F: ttttctctctcgcggccc, R: accacacccctcttatttgtag), *nkx2.2b* (NM_001007782.2, F: TGACTCCACAGATCCACCCT, R: ACAAAAGCGCTCCGTGCC), *sox19a* (NM_130908.2), *tfap2c* (NM_001008576.2, F: tgcggccatgacctcgct, R: tgctggccacgcccactc), *chrd* (NM_130973.3 F: AGCTGCGCGGACAGATAC, R: GGAGACGCCGGTGGGAAAATCCA), *tbxta* (NM_l31162.1), and *sox32* (NM_l31851.1, F: cgaccggatgctccctgacc, R: gcagcaatctggatggaagcagca).

Embryos were fixed and processed as described above in “Immunofluorescence and fluorescence microscopy“. Embryos were incubated with DIG-labeled antisense RNA probes produced using a DIG RNA Labeling Kit (Roche, 11277073910). Probes were detected using anti-digoxigenin-AP Fab fragments (Roche, 11093274910) and developed with BM Purple (Roche, 11442074001). Embryos were fixed in 4% formaldehyde in PBSTx and imaged on a Leica M205 FCA stereoscope.

To quantify the expression patterns of *szl,* and *nkx2.2b,* images taken using consistent parameters were imported into FIJI image J. During quantification, investigators were blinded to the title of the images. To prevent embryo size from confounding our analysis, for the *nxk2.2b* domain size quantification, the anterior-to-posterior length of the domain was divided by the embryo diameter and converted to a percentage. For the *szl* in situ, the FIJI Color Threshold tool was used with consistent settings to measure the area of expression, which was divided by the entire embryo’s area and converted to a percent. Raw data for the in situ quantifications are available in Data S10.

### RNA fluorescence in situ hybridization (RNA-FISH)

Embryos were collected and processed as described above in “Immunofluorescence and fluorescence microscopy“, then RNA FISH was conducted using the Molecular Instruments’ protocol for HCR-Gold RNA FISH. A probe for *tent5ba* was custom-designed and ordered from Molecular Instruments. On day 1, embryos were rehydrated and incubated in probe and probe hybridization buffer at 37°C overnight. On day 2, embryos were washed with probe wash buffer and amplified in 0.06 pmol of hairpins per microliter. On day 3, samples were washed in SSCT, counterstained with SYTOX Green, mounted in ProLong Gold Antifade, and imaged on a Leica Stellaris 5 laser-scanning confocal microscope. Images were acquired on whole-mounted embryos encompassing the entire embryo. To quantify the *tent5ba* HCR in Imaris, a surface was created using the SYTOX Green nuclear signal using consistent parameters. From each surface, the total fluorescent intensity from the *tent5ba* HCR and the surface’s volume in cubic microns were exported. To account for embryo size differences, we divided the sum of the fluorescence by the volume of the embryo. Raw data for HCR FISH quantification is available in Data SlO.

### Lactate measurements

Wildtype or *MZtent5ba*^-42/-42^ embryos were collected after either **1)** 6 hours of normoxic development or 2) 6 hours ofnormoxic development and 18 hours of hypoxic conditions. For each replicate, two embryos per condition were flash-frozen in PCR tubes on dry ice and stored at −80°C. Frozen embryos were processed, and extracts were measured using the Lactate-Glo Assay kit (Promega, 15021) following the manufacturer’s guidelines. For sample processing, the frozen embryos were thawed in 2:1 PBS/inactivation solution (0.6 N HCl) and mixed for 5 minutes. Extracts were neutralized by adding 1 M Tris base (pH 10.7). Embryo lysates, blank wells, and a standard curve oflactate were analyzed according to the kit’s protocol. The luciferase’s bioluminescence was measured using a FlexStation 3 Multi-Mode Microplate Reader (Molecular Devices, Flex3). The background signal, calculated as the average of the blank well’s luminescence, was subtracted from all wells. Lactate concentrations were calculated by fitting the background-subtracted signal to a lactate titration curve. To compare replicates conducted over multiple days, lactate concentrations per embryo were normalized to the experiment’s average normoxic wild-type lactate concentration. Raw data for lactate measurements is available in Data Sl0.

### Nano3p-Seq

Wildtype *(tent5ba^+/+^)* and mutant (MZ*tent5ba*^42/-42^ horionated after either 1) 6 hours of normoxic development or 2) 6 hours of normoxic development and 18 hours of hypoxic conditions. Total RNA was extracted from 25 embryos using the same protocol described in the Bulk RNA sequencing section above. The concentration of total RNA was measured using the RNA Qubit RNA RB Assay Kit (lnvitrogen, Q10210), with readings obtained on the Qubit 2.0 Fluorometer (Invitrogen, Q32866). Given the reported biases that olido(dT) methods can introduce, we opted to perform rRNA depletion to enrich for *mRNA*(*93, 94*). One µg of total RNA per lysate was input into the RiboCop rRNA Depletion Kits for Fish (Lexogen, 241), where the manufacturer’s protocol was followed. The quality and quantity of rRNA-depleted RNA were measured using a High Sensitivity RNA ScreenTape (Agilent, 5067-5579) on an Agilent TapeStation 4200. If necessary, samples were merged and concentrated using a 1.8x SPRI bead cleanup (Beckman Coulter, A63881). Approximately 25 ng ofrRNA-depleted RNA for each library was used for Nano3p-seq, which involved reverse transcription by template switching, complementary DNA annealing, native barcode ligation, and native adapter ligation (Oxford Nanopore Technologies, SQK-NBDl 14-24), all according to the previously published protocol from the Novoa lab(52, 53). Library sizes were estimated using a High Sensitivity RNA ScreenTape, and concentrations were assessed using a Qubit dsDNA HS Assay Kit (Invitrogen, Q32851).

Libraries were sequenced at the UC Davis Genome Center, DNA Technologies and Expression Analysis Core, on a PromethION (Oxford Nanopore Technologies). POD5 files were basecalled using dorado’s (0.7.2) ’super-accurate’ basecalling model (dna_r10.4. l_e8.2_400bps_sup@v5.0.0). Using minimap2 (2.30) and samtools (1.22.1), reads were aligned to the zebrafish genome (danRerl 1). Transcripts were annotated and quantified by PolyTailor’s script “get_transcript_ends.py” and IsoQuant (3.7.1) to the zebrafish annotated transcriptome (V4.3.2.gtt) from the Lawson Lab(67). Poly(A) tail lengths and compositions were estimated using PolyTailor’s script “get_pT.py“. An adjusted poly(A) length was calculated by multiplying the poly(A) length by 1.25 and adding 2, as per PolyTailor’s documentation. Only high-quality reads (passed PolyTailor’s “OK” filter) that mapped to a gene were included in further analysis. To compare our dataset’s accuracy, we exported the source data from Fig. 3G of Begik et al. 2022(52) for 6 hpf embryos and compared the median estimated PAL for each gene (with more than 20 reads) across studies.

Reads were grouped by condition, and adjusted poly(A) tail lengths were compared using a two-sided Mann-Whitney U test, with multiple comparisons adjusted via the Benjamini-Hochberg method. Genes were only evaluated statistically if there were more than 19 reads in each group. Poly(A) tail lengths and differential expression (from either Illumina bulk RNA seq and Nano3P-seq RNA seq) were graphed using matplotlib (3.10.0) and seaborn (0.13.2).

Regression lines were calculated using a simple least-squares regression. Correlation metrics were calculated using either a Pearson’s and/or Spearman’s correlation test, indicated in the figures and/or captions. Differential gene expression using Nano3p-seq libraries was conducted as described above using DESeq2. Nano3p-seq summary data, differential polyadenylation, and differential expression are available in Data S8, S10, and S11.

### Statistical analysis

Statistical analysis was performed in GraphPad Prism (10.4.2), R (4.3.1), and Python (3.12.12). Particular statistical tests used in this study are specified in the figure captions and/or methods. In all figures, statistical significance is indicated as*= P≤0.05, ** = P≤0.01, *** = P≤0.001, **** = P≤0.0001.

## Data Sl. (Separate file)

Single cell summary statistics and cNMF gene expression programs. Single-cell quality metrics. All gene expression programs (GEPs) that were identified using consensus non-negative matrix factorization (cNMF). The 2000 genes associated with each GEP are ordered by their level of association.

## Data S2. (Separate file)

Differential gene expression for all single cell cluster comparisons between normoxia 6 hpf and hypoxia 2hrs. The results of pseudobulked differential gene expression from single cell cluster comparisons between normoxic and hypoxic conditions. Columns are paired by cluster, one depicting the log2(fold change) and the other the adjusted P-value.

## Data S3. (Separate file)

Bulk RNA-seq batch information and differential gene expression. Batch information for various bulk RNA-seq samples. The results of differential gene expression tests from bulk RNA seq experiments across many conditions including: **1)** Reoxy 4hrs vs Normoxia 10 hpf 2) Hypoxia 2 hrs vs Normoxia 6 and 8 hpf 3) Hypoxia 18hrs vs Normoxia 6 and 8 hpf 4) KCN 18 hrs vs Hypoxia 18 hrs 5) Hypoxia 18 hrs *tent5ba*^-42/-42^ vs Hypoxia 18 hrs *tent5ba*^+/+^ 6) Reoxy 4 hrs *tent5ba*^-42/-42^ vs Reoxy 4 hrs *tent5ba^+/+^*.

## Data S4. (Separate file)

Differential accessibility from bulk ATAC-seq. Differential accessibility between normoxia 8 hpf and hypoxia 18hrs. Table includes peak name, peak location (genomic coordinates), concentrations by group, log2(fold change) for each peak, P-value, false discovery rate (FDR), and the closest gene ([-2000, +500] bp from TSS).

## Data S5. (Separate file)

Gene set enrichment analysis (GSEA) and Overrepresentation analysis (ORA) results. Aggregated tables for GSEA and ORA tables.

## Data S6. (Separate file)

Dormancy Enriched Expression Program associated genes. Aggregated ranked gene lists from cNMF (GEP 5), scRNA-seq (common differentially expressed (log2FC<l, P-adj<0.05) genes across cluster, ordered by frequency and then differential expression magnitude), bulk RNA-seq acute (hypoxia 2hrs vs normoxia 6 and 8 hpf, log2FC>**1,** P-adj<0.05, ordered by log2FC magnitude), bulk RNA-seq prolonged (hypoxia 18 hrs vs normoxia 6 and 8 hpf, log2FC>l, P-adj<0.05, ordered by log2FC magnitude), bulk RNA-seq KCN paused (KCN 18 hrs vs normoxia 6 hpf, log2FC>l, P-adj<0.05, ordered by log2FC magnitude), and bulk ATAC-seq (hypoxia 18 hrs vs normoxia 8 hpf, log2FC>1, P-adj<0.05, ordered by log2FC magnitude, highest occurrence only). Genes within each column were assigned a score based on their normalized column order, with an absence resulting in the lowest score. Genes were then ordered by their ’DEEP’ score in the last column.

## Data S7. (Separate file)

*tent5ba* dependent DEEP transcripts. Contrasted differential gene expression results from hypoxia 18 hrs *tent5ba*^+/+^ to normoxia 6 hpf *tent5ba*^+/+^ (log2FC2“..0.5, P-adj .’.S0.05) and hypoxia 18 hrs *tentSba*^-42/-42-42/-42^ *tent5ba*^+/+^ (log2FC.’.S-0.5, P-adj.’.S0.05). Transcripts that fit these criteria have a ’Yes’ flag in the column *tent5ba* dependent DEEP transcripts, otherwise they remain blank.

## Data S8. (Separate file)

Nano3p-seq summary and differential polyadenylation. Summary statistics ofNano3p-seq, median and average poly(A) tail lengths per group, and differential polyadenylation.

## Data S9. (Separate file)

Exon intron changes across conditions. Per-gene statistical results (Δlog2FC and FDR) quantifying the difference between exonic and intronic fold-changes, used as a measure of post-transcriptional regulation across conditions.

## Data Sl0. (Separate file)

Raw data and statistics. Raw data and statistics for epiboly quantification, morphogen gradients, larval survival after pausing, hypoxia tolerance, single cell cluster proportions, Euclidean PC distances, lactate measurements, exon intron mapping ratios, *tent5ba* HCR quantification, *tent5ba* dorsalization frequencies, reoxy pSmadl/5/9 quantification, and in situ hybridization quantifications.

## Data Sll. (Separate file)

Nano3p-seq differential gene expression. The results of a differential gene expression test between normoxia 6 hpf *tent5ba^+I+^* and normoxia 6 hpf *MZtentSba*^-42/-42-42/-42^ embryos.

---

**Fig. S1.**
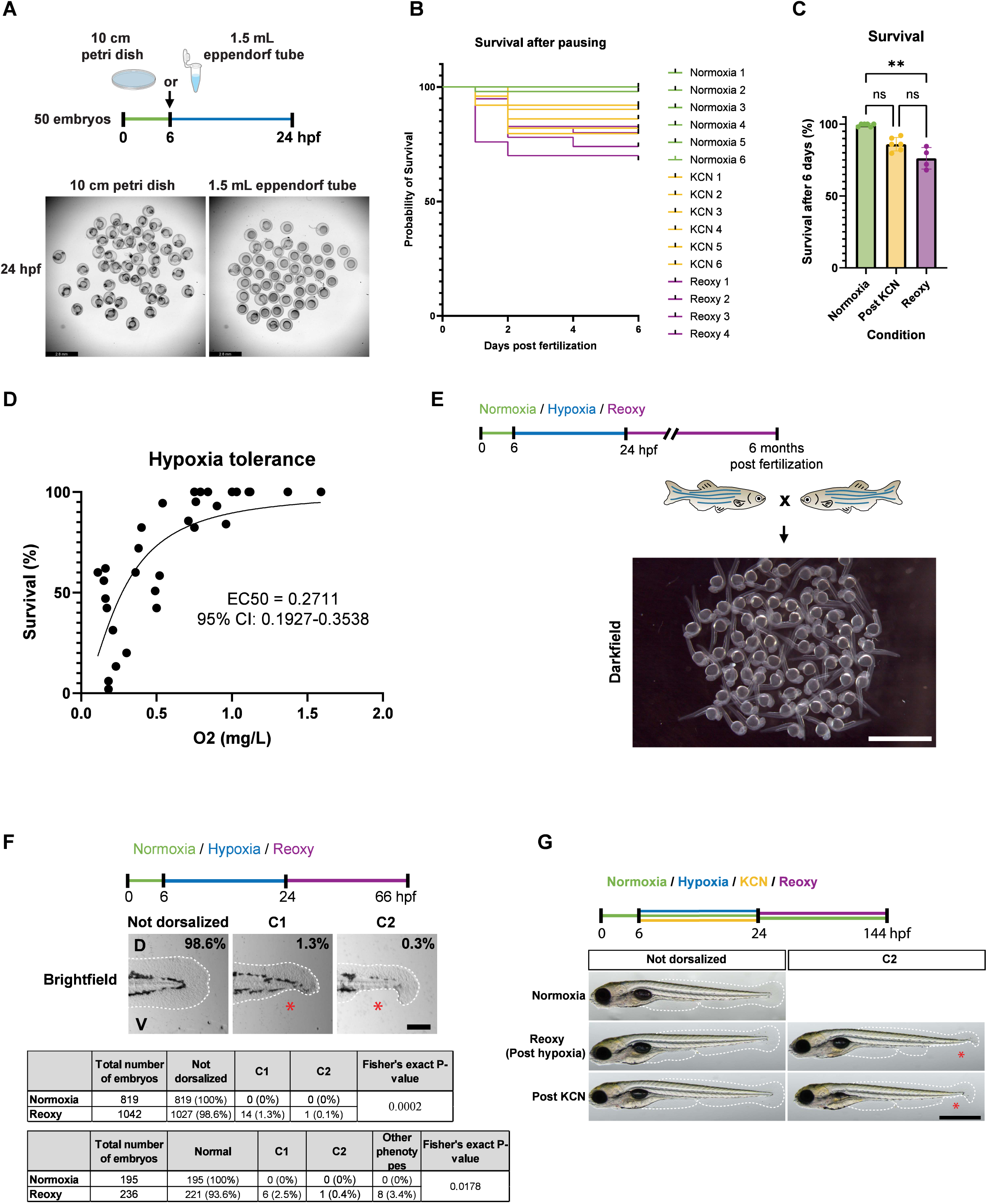
Impacts of hypoxia-induced pausing on survival, fertility, and developmental defects. **(A)** Brightfield images of ∼50 embryos after either standard or crowded conditions. Scale, 2.8 mm. **(B, C)** Survival of cohorts (∼50 embryos) exposed at 6 hpf to 18 hrs of normoxia, KCN, or hypoxia. (C) Quantification of survival in bar plots (mean±SD). Kruskal-Wallis and Dunn’s test. **(D)** Oxygen concentration (measured at end) and survival of reoxy 24 hrs embryos. Dots (∼50 embryos), n=38. Data also in Fig. 4A and S8C. **(E)** Darkfield image of an embryo clutch produced from zebrafish previously paused for 18 hrs at shield stage. Scale, 2.9 mm. **(F)** Images and table of observed dorsalization phenotypes. Whole-mount images of 6 hpf embryos exposed to 18 hrs of hypoxia and then cultured in normoxia for ∼42 hrs. Scale, 173 µm. ’D’=Dorsal and ’V’=Ventral. (F, G) Red asterisks indicate reduction/lack of ventral tail fin. **(G)** Brightfield images of embryos exposed to 18 hrs of normoxia, hypoxia, or KCN at shield and then normoxia for ∼120 hrs. Scale, 832 µm. In all figures, (P or adjusted P-values): * = P≤0.05, ** = P≤0.01, *** = P≤0.001, **** = P≤0.0001 (Methods).

**Fig. S2.**
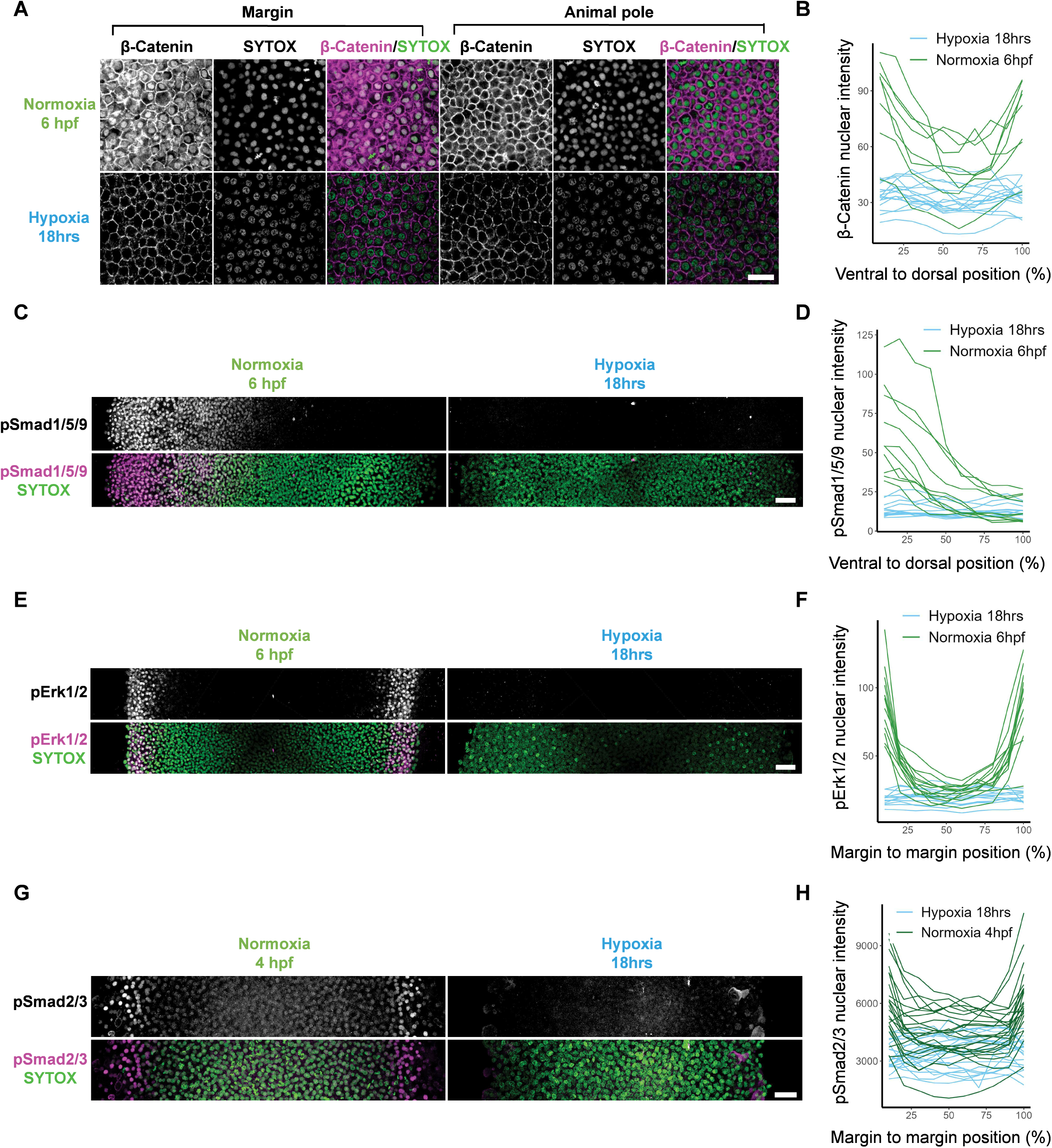
Cessation of morphogen gradients during pausing. **(A)** Representative immunofluorescent z-planes of β-Catenin and SYTOX Green staining in embryos at 6 hpf or those that at 6 hpf had been exposed to 18 hrs of hypoxia. These images are extensions of Fig. 1D. Scale, 26 µm. **(B, D, F, H)** Individual embryo quantifications of immunofluorescence stains for the nuclear intensity of β-Catenin, pSmadl/5/9, pErkl/2, or pSmad2/3 across the designated embryonic axis. (B, D, F) either 6 hpf or 6 hpf exposed to 18 hrs of hypoxia. (H) either 4 hpf or 4 hpf exposed to 18 hrs of hypoxia. These plots are the individual embryo traces that comprise Fig. lE-H. For B, normoxia n=8, hypoxia n=16. For D, normoxia n=l0, hypoxia n=13. For F, normoxia: n=23; hypoxia: n=21. For H, normoxia n=13, hypoxia n=12. **(C, E, G)** Representative immunofluorescent maximum projections across embryos. (C, E) Embryos at 6 hpf or those that at 6 hpf had been exposed to 18 hrs of hypoxia, (G) embryos at 4 hpf or those that at 4 hpf had been exposed to 18 hrs of hypoxia. Scale bars, (C) 57.4 µm, (E) 47.2 µm, (G) 37.1 µm.

**Fig. S3.**
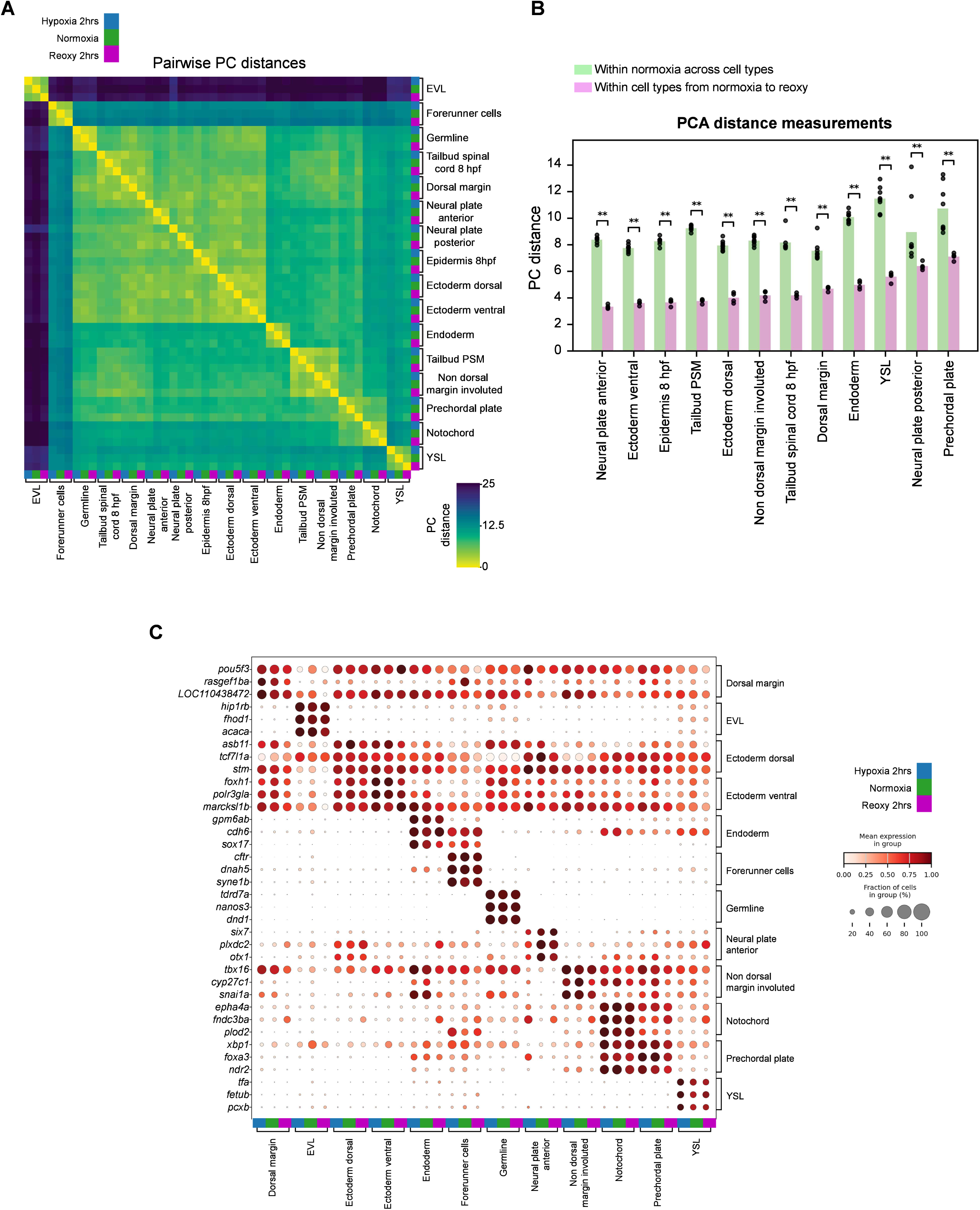
Cell identity retention during pausing. **(A)** Heatmap of the pairwise PC Euclidean distances between cells binned by their cell type and condition. Conditions are indicated by colors, where blue is hypoxia 2hrs, green is normoxia 6 hpf, and purple is reoxy 2 hrs. **(B)** Euclidean distance measurements of PC space. In green, the average distance between the specified cell type in normoxia and all other cell types within normoxia. In purple, the distance between a cell type in normoxia 6/8 hpf and reoxy 2 hrs. The 12 most abundant cell types were assayed. Each dot represents a replicate’s value, P-values from the Mann-Whitney U test. **(C)** Dotplot of the top 3 non-redundant marker genes for each cell type in normoxia. The mean expression and fraction expressing these marker genes are ordered by cell type annotation and condition, hypoxia 2 hrs in blue, normoxia 6 hpf in green, and reoxy 2 hrs in purple. In all figures, (P or adjusted P-values): * = P≤0.05, ** = P≤0.01, *** = P≤0.001, **** = P≤0.0001 (Methods).

**Fig. S4.**
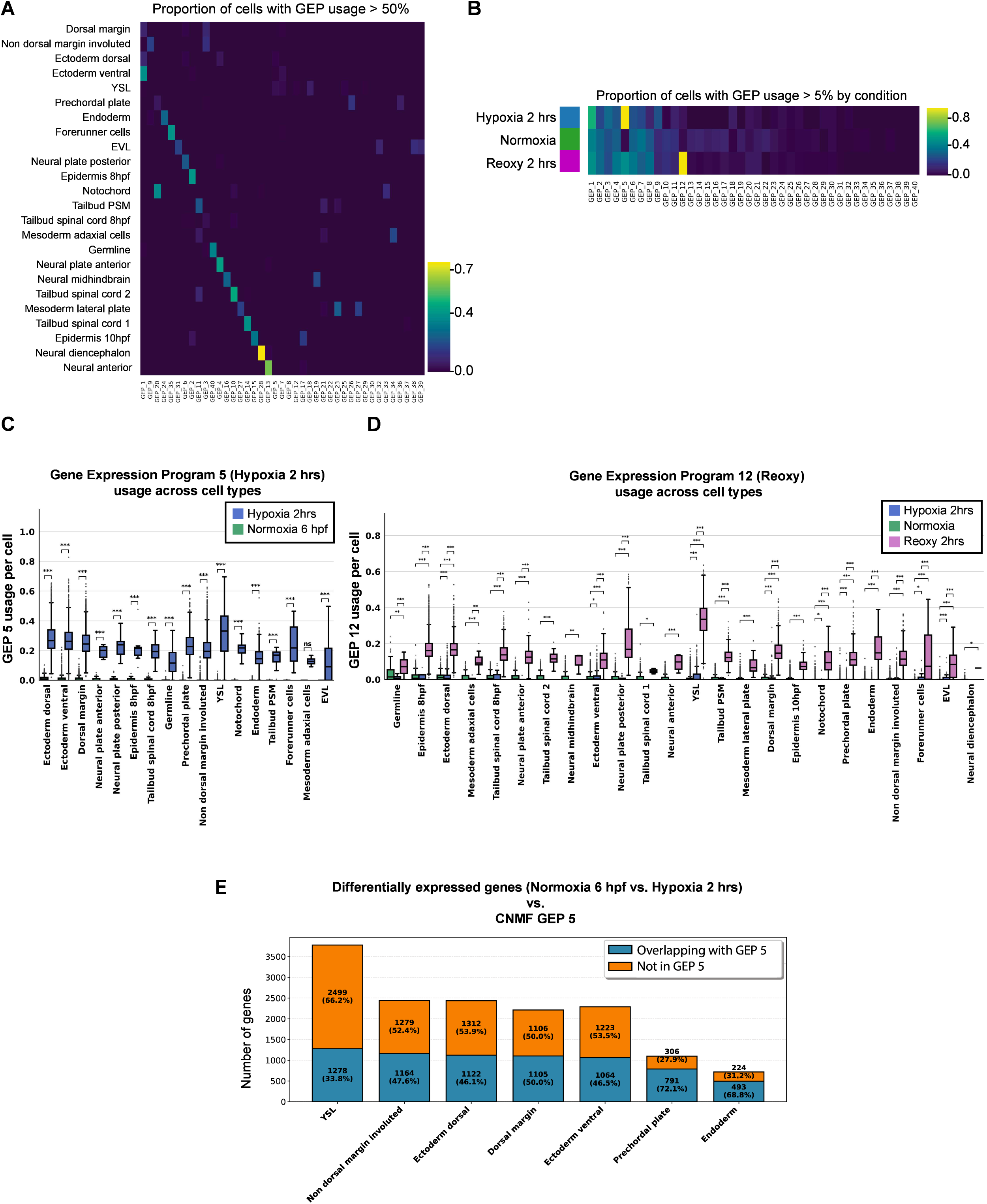
DEEP is orthogonal to cell identity programs. **(A)** Heatmap of the cNMF gene expression program (GEP) usage scores across cell type annotations. The colormap intensity corresponds to the proportion of cells within a cell type annotation that have a GEP usage score greater than 50%. This plot comprises cells from normoxia, hypoxia, and reoxy. **(B)** Heatmap of the cNMF GEP usage scores across conditions. The intensity of the colormap is proportional to the percentage of cells in a condition with a GEP usage score greater than 5%. This plot comprises cells from normoxia, hypoxia, and reoxy. **(C, D)** Boxplots showing interquartiles, range, and outliers of the cNMF-derived Gene Expression Programs, (C) GEP 5, (D) GEP 12, usage score per cell, separated by condition. In (D) The condition normoxia includes cells from 6, 8, and 10 hpf. Mann Whitney U test, in (D) BH corrected. **(E)** Stacked bar plot comparing genes up-regulated in hypoxia 2 hrs compared to normoxia 6 hpf (P-adj<0.05, log2FC<-0.5, from psuedobulk via pyDEseq2) with the top 2000 genes in the cNMF-derived GEP5. If these genes were identical, they were labeled as “Overlapping with GEP 5”. See Data SI, S2. In all figures, (P or adjusted P-values): * = P≤0.05, ** = P≤0.01, *** = P≤0.001, **** = P≤0.0001 (Methods).

**Fig. S5.**
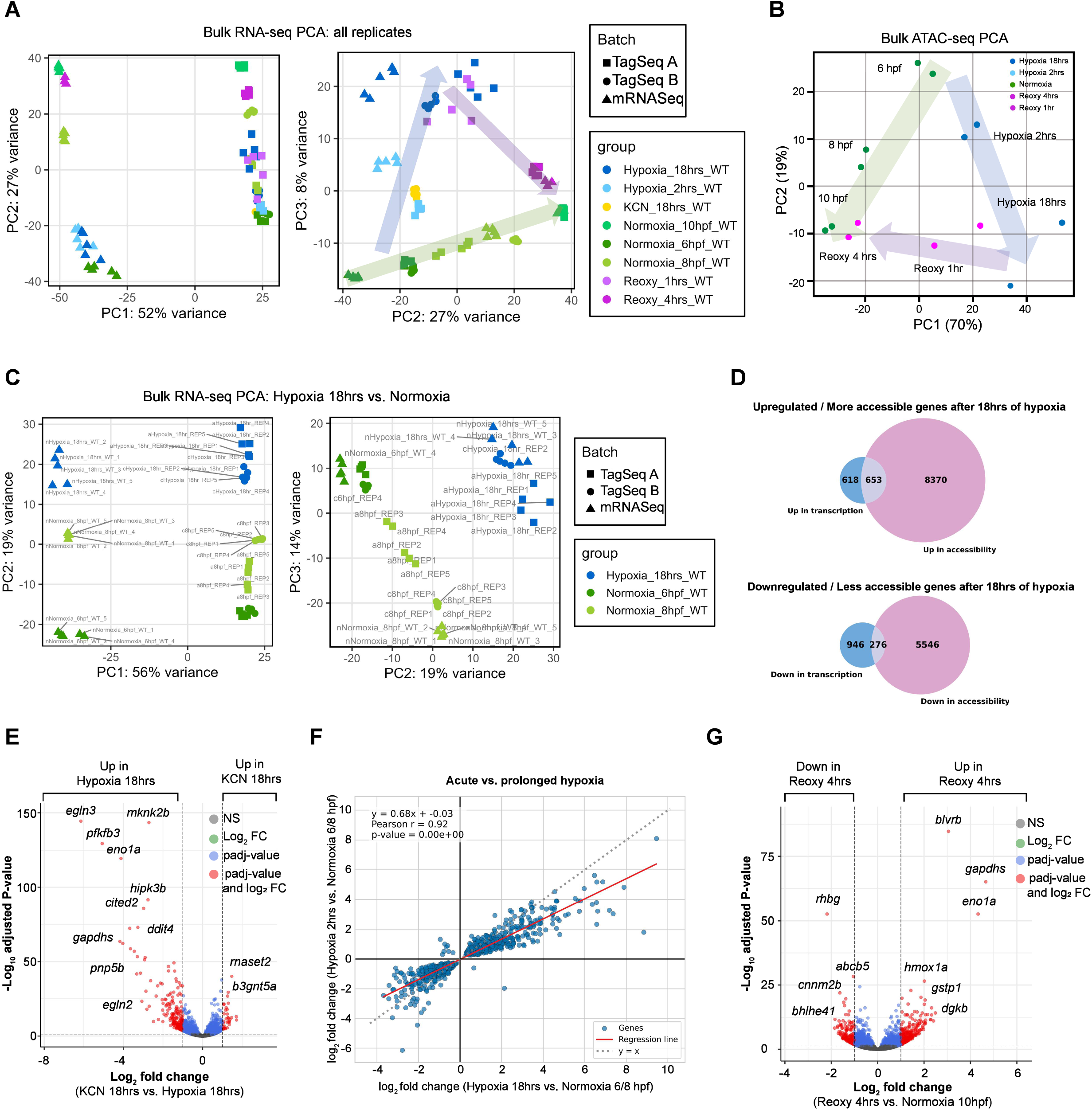
DEEP across modality, pausing paradigm, technology, and time. **(A)** PCA for three batches of bulk RNA-seq datasets, PCl vs PC2 and PC2 vs PC3. Arrows added manually. **(B)** PCA of bulk ATAC-seq dataset. Arrows manually added. **(C)** PCA of samples across all batches used for the normoxic-to-hypoxic comparison. **(D)** Venn diagram comparing the differential expression to differential accessibility after 18 hrs of hypoxia. Differentially expressed genes (P-adj<0.01, llog2FCl>l) from hypoxia 18 hrs and normoxia 6 and 8 hpf samples were contrasted with differentially accessible peaks (FDR<0.01, llog2FCl>l) from hypoxia 18 hrs and normoxia 8 hpf samples. See Data S3, S4. **(E)** Volcano plot of differentially expressed genes between 6 hpf embryos exposed to either 18 hrs of KCN or hypoxia. Well-separated points were manually labeled, see Data S3. **(F)** Acute (2 hrs) vs prolonged (18 hrs) hypoxia, only genes P-adj≤0.05. In red, linear least-squares regression line. Pearson’s correlation and Wald’s. See Data S3. **(G)** Volcano plot of differentially expressed genes between reoxy 4 hrs and normoxia 10 hpf embryos. Well-separated points manually labeled. See Data S3.

**Fig. S6.**
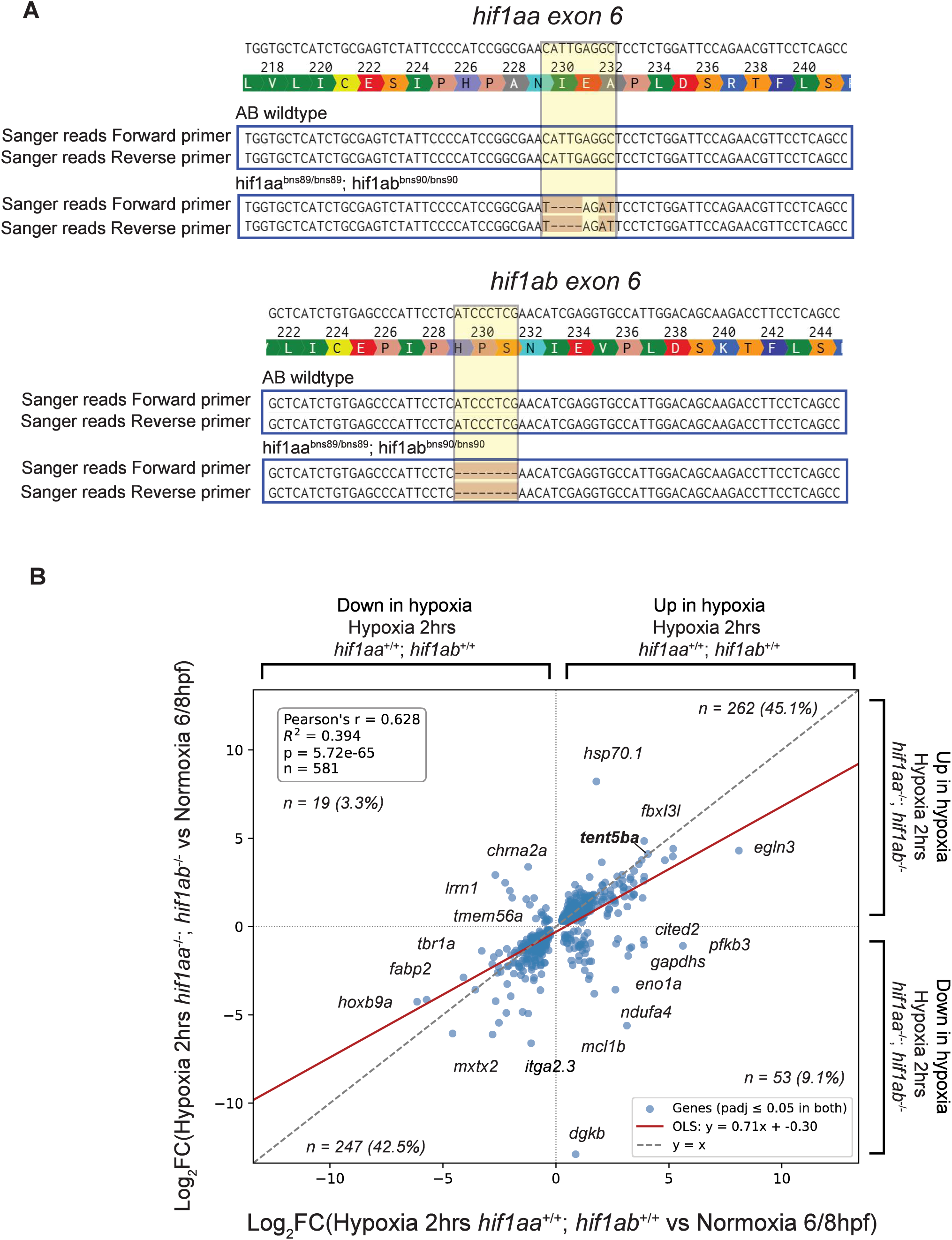
***tent5ba* expression in hypoxia is not dependent on *hifl aa* and *hifl ab* (A)** Aligned Sanger sequencing reads of PCR products from the *hifl aa* and *hifl ab* loci from AB wildtype and *hifl aa^bns89/bns89^;hiflbns90^/^bns90* (also referred to *hifl aa^-/-^;hifl ab^-I-^)* fin clippings. **(B)** Comparison of differential gene expression between AB wildtype embryos in normoxia and either AB wildtype or *hifl aa^-/-^;hifl ab^-/-^* embryos exposed to 2 hrs of hypoxia. Only genes that are differentially (P-adj≤S0.05) expressed in both comparisons are graphed. Linear least-squares regression line in red. Well-separated points were annotated with gene names manually.

**Fig. S7.**
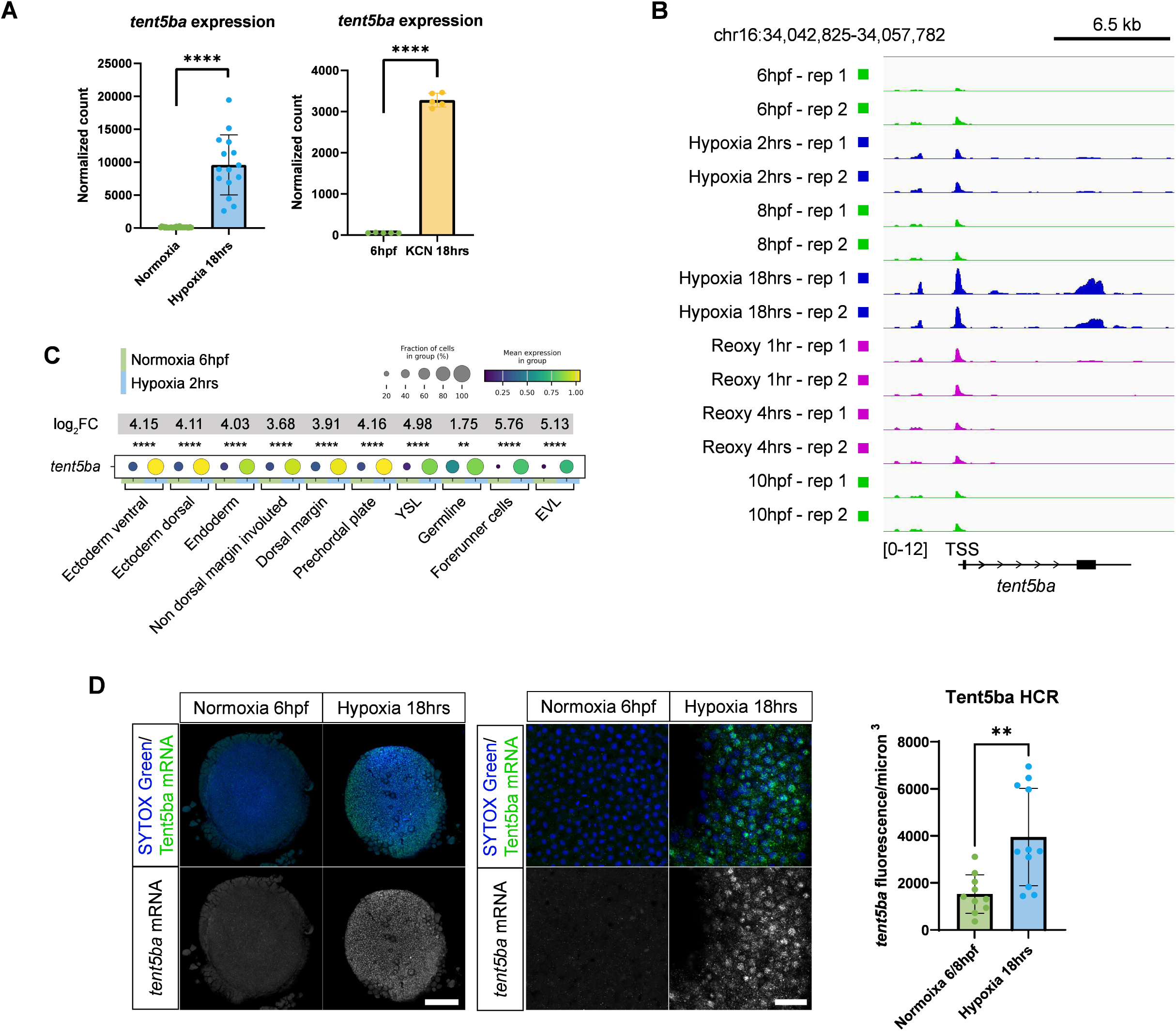
***tent5ba* expression is increased across the embryo upon hypoxic exposure (A)** *tent5ba* expression from bulk RNA sequencing, in 6 hpf embryos in normoxia or after they had been exposed to 18 hrs of either hypoxia or KCN. Adjusted P-values from DESeq2, see Data S3. **(B)** *tent5ba* locus across the specified conditions overlaid with ATAC-seq accessibility tracks, see Data S4. **(C)** Dotplot of *tent5ba* expression from scRNA-seq across cell types in normoxic 6 hpf embryos and 6 hpf embryos exposed to 2 hrs of hypoxia. Log2 fold changes and adjusted P-values from pyDESeq2, see Data S2. **(D)** *tent5ba* expression from HCR RNA FISH in embryos at 6 hpf in normoxia and after 18 hrs of hypoxia. The first panel shows representative low-magnification maximum projections (∼100 µm) of animal poles from whole-mounted embryos; scale bar: 154 µm. The second panel includes higher-magnification images of the animal poles of whole-mounted embryos; scale bar: 30 µm. Quantification (mean±SD) of the total *tent5ba* HCR fluorescence within and relative to the volume of the embryo within the higher-magnification z-stacks (normoxia: n=lO, hypoxia: n=l 1). P-values from a Mann-Whitney U test. In all figures, (P or adjusted P-values): * = P≤0.05, ** = P≤0.01, *** = P≤0.001, **** = P≤0.0001 (Methods).

**Fig. S8.**
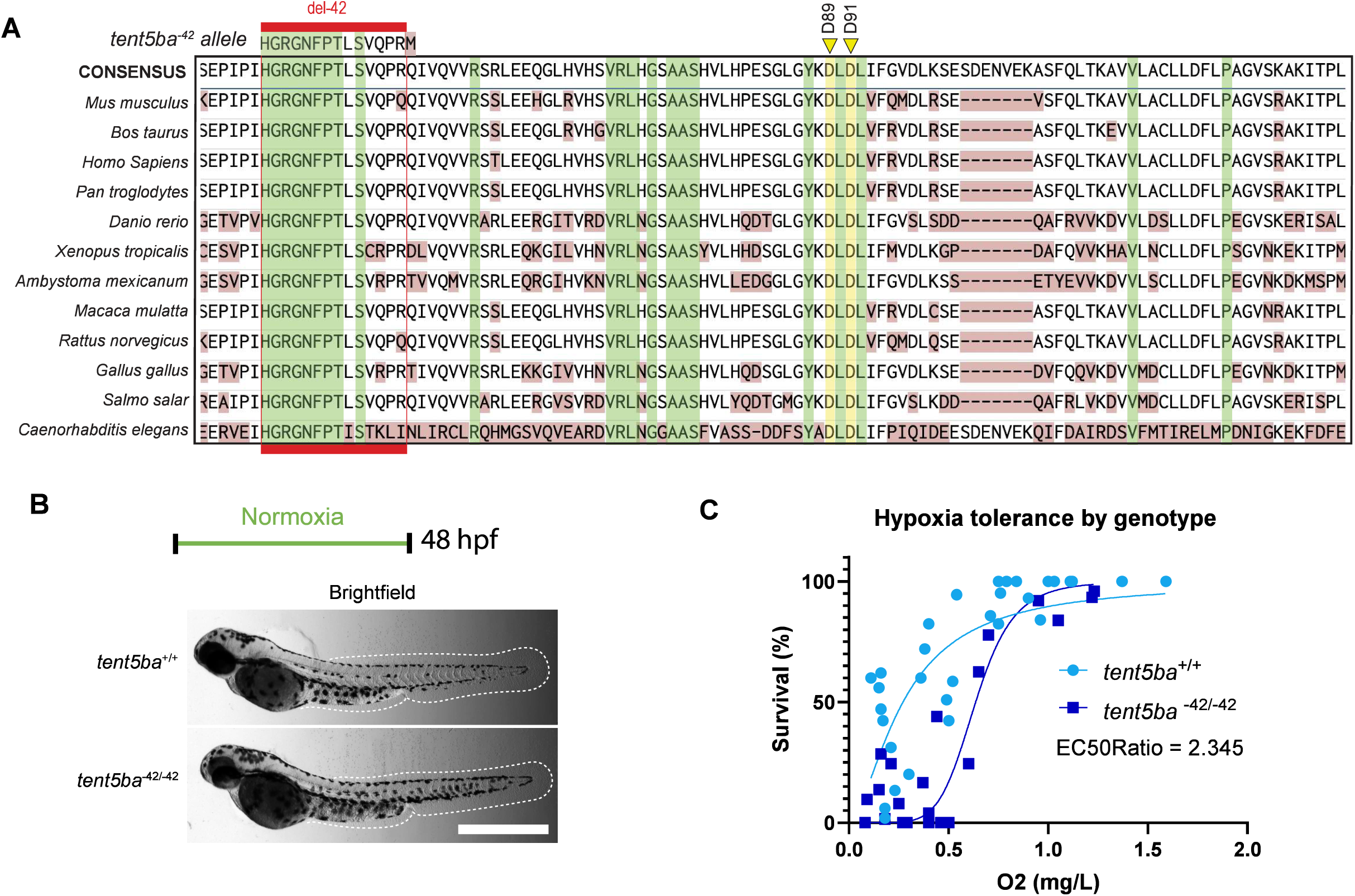
***tentSba-^42^* allele and its role in hypoxia tolerance (A)** Multiple sequence alignment of a region (Danio rerio: G29-Ll31) of the N-terminal catalytic domain region of Tent5b proteins. *tent5ba-^42^* row indicates the amino acid sequence corresponding to its indel, deletions within the red box (del-42) plus a Q49M. Yellow indicates conserved catalytic aspartic acid residues putatively responsible for Tent5b’s nucleotidyltransferase activity, corresponding to D124 and D126 in human, respectively. Green represents amino acid conservation and rust represents divergence. **(B)** Representative images of normoxic *tent5ba^+/+^and MZtent5ba*^-42/-42^. Scale, 794 µm. **(C)** Survival(%) of *tent5ba*^+/+^(n=31) and *MZtentSba*^-42/-42^ (n=l 7) reoxy 24 hrs. Survival vs oxygen concentration (at end). EC50 via least-squares regression nonlinear curve. Dots represent replicates (∼50 embryos each). Data also in Fig. SID and 4A.

**Fig. S9.**
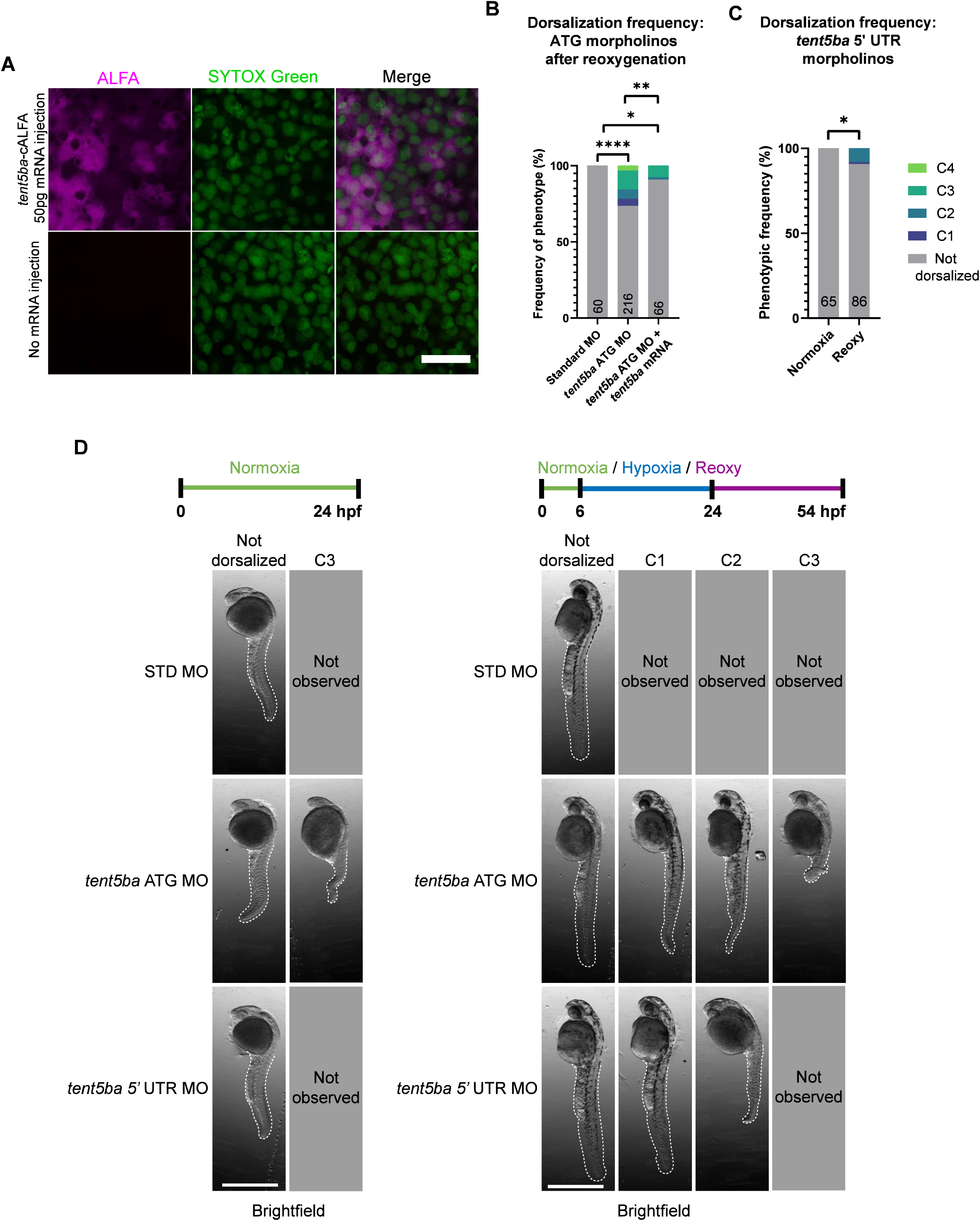
***tent5ba* overexpression and knockdown (A)** Representative immunofluorescence z-slices of anti-ALFA staining in 6 hpf embryos. *tent5ba^+I+^* embryos were injected with mRNA encoding *tent5ba* transcript fused to an ALFA tag on its C-terminal end prior to the stop codon. Scale bar, 110 µm. **(B, C)** Prevalence of dorsalized phenotypes observed in *tent5ba*^+/+^ embryos injected with (B) translation-blocking morpholinos targeting the human beta globin intron (standard), *tent5ba* start codon (ATG), and/or *tent5ba-*ALFA mRNA after reoxygenation or (C) translation-blocking morpholinos targeting the 5’ UTR of *tent5ba* in normoxia and after reoxygenation. P-values from a Fisher’s exact test (n in plots). **(C)** Representative images of dorsalized or normal phenotypes in normoxic and reoxygenated conditions after injections of STD MO *tent5ba,* ATG MO, or *tent5ba* 5’UTR MO Scales bars, 576 µm. In all figures, (P or adjusted P-values): * = P≤0.05, ** = P≤0.01, *** = P≤0.001, **** = P≤0.0001 (Methods).

**Fig. S10.**
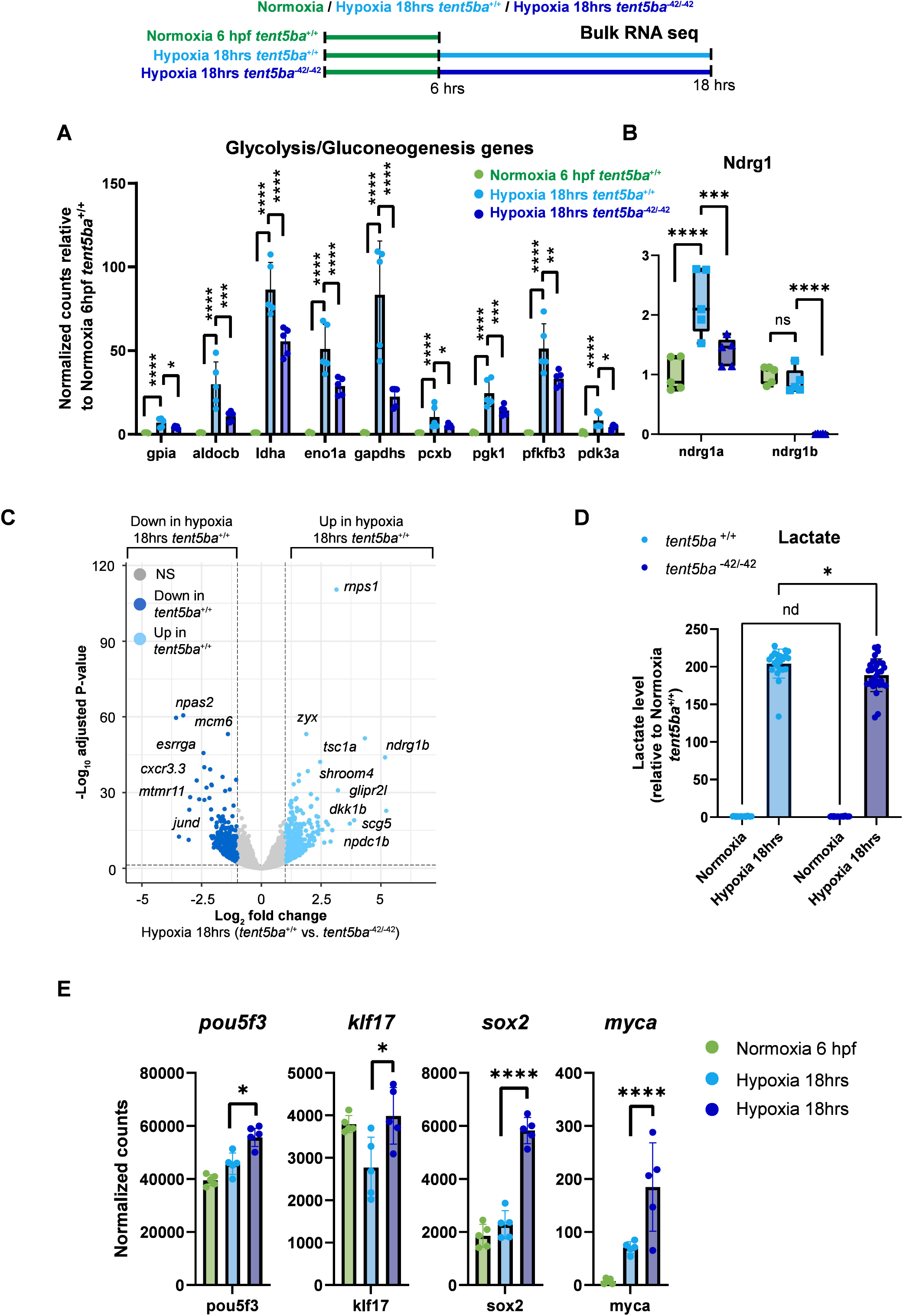
Paused gene expression in *MZtentSba-^42/^-^42^* mutants. **(A)** Schematic bulk RNA-seq. Bar plots of glycolytic/gluconeogenic-related genes from mRNASeq. (A, B) Normalized counts relative to normoxia 6 hpf *tent5ba+I+_*Pairwise P-adj from DESeq2, see Data S3. **(B)** *ndrgla* and *ndrglb* expression from mRNAseq. **(C)** Volcano plot comparing *tent5ba*^+/+^ and *MZtent5ba-^421^_-_^42^* hypoxia 18hrs, differentially expressed genes (P-adj<0.05, llog2FCI> **1).** Well-separated points labeled, see Data S3. **(D)** Lactate levels measured in 6 hpf *tent5ba*^+/+^ (normoxia: n=24, hypoxia: n=22) and *MZtent5ba-^421^_-_^42^* (normoxia: n=24, hypoxia: n=30) after 18 hrs of hypoxia. Mann-Whitney U and two-stage set-up (Benjamini, Krieger, and Yekutieli). **(E)** Bar plots of epiblast marker genes from mRNASeq. Pairwise P-adj from DESeq2, see Data S3. In all figures, (P or adjusted P-values): * = P≤0.05, ** = P≤0.01, *** = P≤0.001, **** = P≤0.0001 (Methods).

**Fig. S11.**
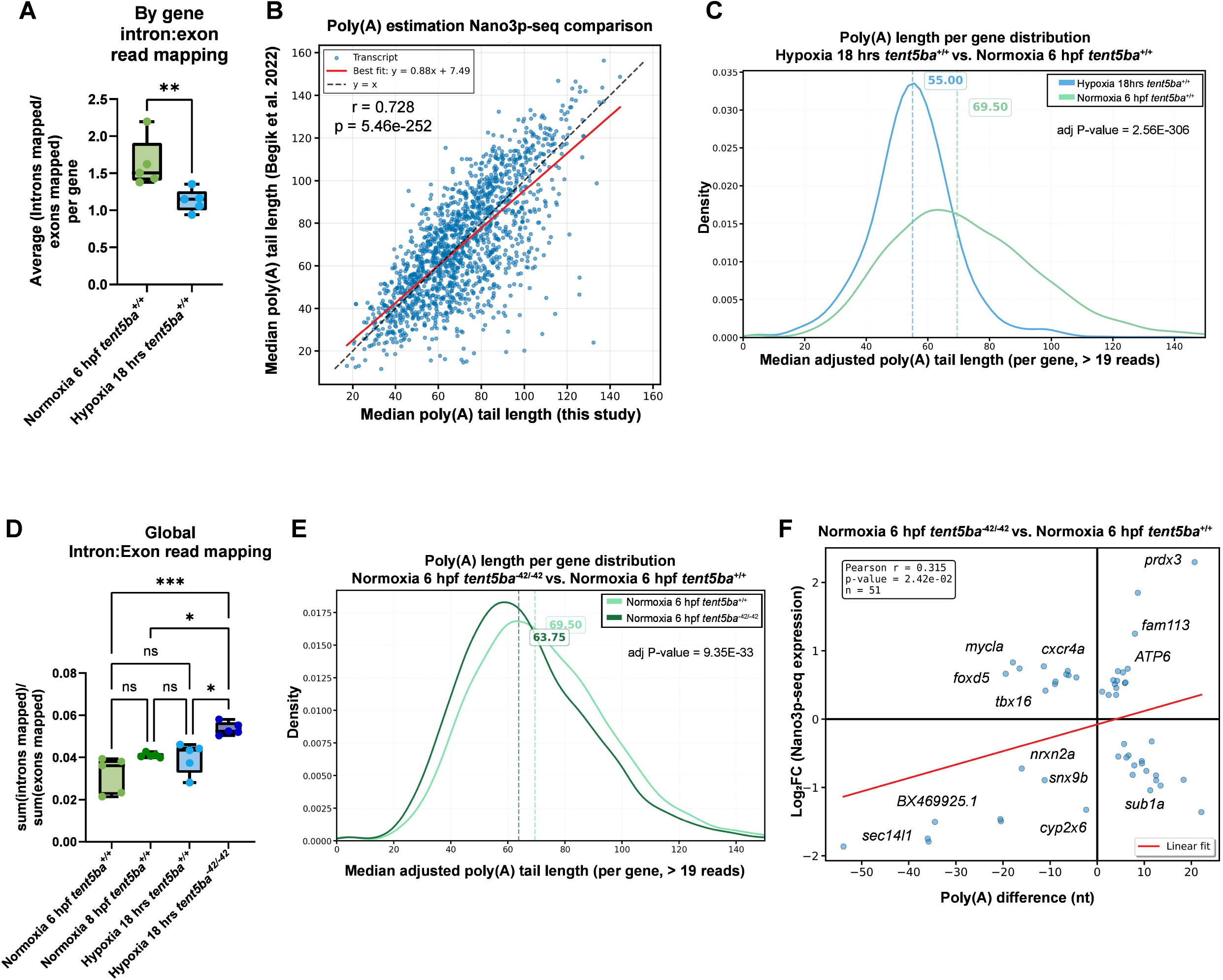
Post-transcriptional dynamics of dormancy and the role of *tentSba*. **(A)** Box (interquartiles) and whisker (min and max) plots of the average intron:exon read mapping ratios per gene. Kruskal-Wallis tests and Tukey method. **(B)** PAL estimations from this study and Begik et al. 2022. Median estimated PAL of transcripts (reads>20 per gene) in normoxic 6 hpf embryos from each dataset. Pearson’s correlation test. Linear least-squares regression in red. **(C, E)** Kernel density estimates of PALs from transcripts with more than 19 reads. Dotted lines are PAL medians from each condition. Plots only include data between 0 and 150 nucleotides. Adjusted P-value from Mann Whitney, **BH** corrected. See also Data S8. (C) Standard deviation for normoxia = 25 nt and hypoxia 18 hrs = 18.1 nt. **(D)** Box (interquartiles) and whisker (min and max) plots of intron:exon read mapping ratios for all transcripts from rnRNASeq. Kruskal-Wallis tests and Tukey method. **(F)** Differential (both P-adj≤0.05) expression (from Nano3p-Seq) vs polyadenylation of 6 hpf *MZtent5ba-^421^-^42^* embryos in normoxia and 6 hpf *tent5ba*^+/+^*embryos* in normoxia, see Data S8, S11. In all figures, **(P** or adjusted P-values): * = P≤0.05, ** = P≤0.01, *** = P≤0.001, **** = P≤0.0001 (Methods).

## References and Notes

1. C. H. Waddington, Canalization of development and genetic assimilation of acquired characters. Nature 183, 1654–1655 (1959).

2. J. Hermisson, G. P. Wagner, The population genetic theory of hidden variation and genetic robustness. Genetics 168, 2271–2284 (2004).

3. H. ö. özgtildez, A. Bulut-Karslioglu, Dormancy, Quiescence, and Diapause: Savings Accounts for Life. Annual Review of Cell and Developmental Biology 40, 25--49 (2024).

4. K. D. Webster, J. T. Lennon, Dormancy in the origin, evolution and persistence oflife on Earth. Proc. Biol. Sci. 292, 20242035 (2025).

5. R. B. Crawford, C. E. Wilde Jr, Cellular differentiation in the anamniota. II. Oxygen dependency and energetics requirements during early development of teleosts and urodeles. Exp. Cell Res. 44, 453--470 (1966).

6. P.A. Padilla, M. L. Ladage, Suspended animation, diapause and quiescence: arresting the cell cycle in C.elegans. Cell Cycle 11, 1672–1679 (2012).

7. B. A. Mendelsohn, J. D. Gitlin, Coordination of development and metabolism in the pre-midblastula transition zebrafish embryo. Dev. Dyn. 237, 1789–1798 (2008).

8. B. A. Mendelsohn, B. L. Kassebaum, J. D. Gitlin, The zebrafish embryo as a dynamic model of anoxia tolerance. Dev. Dyn. 237, 1780–1788 (2008).

9. C. B. Kimmel, W.W. Ballard, S. R. Kimmel, B. Ullmann, T. F. Schilling, Stages of embryonic development of the zebrafish. Dev. Dyn. 203, 253–310 (1995).

10. M. C. Mullins, M. Hammerschmidt, D. A. Kane, J. Odenthal, M. Brand, F. J. van Eeden, M. Furutani-Seiki, M. Granato, P. Haffter, C. P. Heisenberg, Y. J. Jiang, R. N. Kelsh, C. Nilsslein-Volhard, Genes establishing dorsoventral pattern formation in the zebrafish embryo: the ventral specifying genes. Development 123, 81–93 (1996).

11. C. S. Hill, Establishment and interpretation of NODAL and BMP signaling gradients in early vertebrate development. Curr. Top. Dev. Biol. 149, 311–340 (2022).

12. M. L. Concha, Zebrafish gastrulation. Annu. Rev. Cell Dev. Biol. 41, 89–134 (2025).

13. A. F. Schier, W. S. Talbot, Molecular genetics of axis formation in zebrafish. Annu. Rev. Genet. 39, 561–613 (2005).

14. P. Haffter, M. Granato, M. Brand, M. C. Mullins, M. Hammerschmidt, D. A. Kane, J. Odenthal, F. J. van Eeden, Y. J. Jiang, C. P. Heisenberg, R. N. Kelsh, M. Furutani-Seiki, E. Vogelsang, D. Beuchle, U. Schach, C. Fabian, C. Nilsslein-Volhard, The identification of genes with unique and essential functions in the development of the zebrafish, Danio rerio. Development 123, 1–36 (1996).

15. N. A. Aponte-Santiago, Y. Su, D. E. Wagner, ZMAP: A single-cell meta-atlas of zebrafish embryonic development reveals a consensus hierarchy of cell identities, bioRxivorg (2026)p. 2026.03.23.713599.

16. D. Kotliar, A. Veres, M.A. Nagy, S. Tabrizi, E. Hodis, D. A. Melton, P. C. Sabeti, Identifying gene expression programs of cell-type identity and cellular activity with single-cell RNA-Seq. Elife 8 (2019).

17. N. A. Sweet, C.-K. Hu, Using diapause as a platform to understand the biology of dormancy. Open Biol. 15, 250104 (2025).

18. J.C. Fenelon, New insights into how to induce and maintain embryonic diapause in the blastocyst. Curr. Opin. Genet. Dev. 86, 102192 (2024).

19. S. Easwaran, D. J. Montell, The molecular mechanisms of diapause and diapause-like reversible arrest. Biochem. Soc. Trans. 51, 1847–1856 (2023).

20. V. Liudkovska, P. S. Krawczyk, A. Brouze, N. Guminska, T. Wegierski, D. Cysewski, Z. Mackiewicz, J. J. Ewbank, K. Drabikowski, S. Mroczek, A. Dziembowski, TENTS cytoplasmic noncanonical poly(A) polymerases regulate the innate immune response in animals. Sci. Adv. 8, eadd9468 (2022).

21. J.-L. Hu, H. Liang, H. Zhang, M.-Z. Yang, W. Sun, P. Zhang, L. Luo, J.-X. Feng, H. Bai, F. Liu, T. Zhang, J.-Y. Yang, Q. Gao, Y. Long, X.-Y. Ma, Y. Chen, Q. Zhong, B. Yu, S. Liao, Y. Wang, Y. Zhao, M.-S. Zeng, N. Cao, J. Wang, W. Chen, H.-T. Yang, S. Gao, FAM46B is a prokaryotic-like cytoplasmic poly(A) polymerase essential in human embryonic stem cells. Nucleic Acids Res. 48, 2733–2748 (2020).

22. K. Kuchta, A. Muszewska, L. Knizewski, K. Steczkiewicz, L. S. Wyrwicz, K. Pawlowski, L. Rychlewski, K. Ginalski, FAM46 proteins are novel eukaryotic non-canonical poly(A) polymerases. Nucleic Acids Res. 44, 3534–3548 (2016).

23. S. Mroczek, J. Chlebowska, T. M. Kulinski, O. Gewartowska, J. Gruchota, D. Cysewski, V. Liudkovska, E. Borsuk, D. Nowis, A. Dziembowski, The non-canonical poly(A) polymerase FAM46C acts as an onco-suppressor in multiple myeloma. Nat. Commun. 8,619 (2017).

24. N. J. Proudfoot, A. Purger, M. J. Dye, Integrating mRNA Processing with Transcription. Cell 108, 501–512 (2002).

25. D. Lacidogna, S. Pennacchio, E. Milan, TENT5/FAM46: An enigmatic family of secretory tuners. Traffic 26, e70011 (2025).

26. C. Fucci, M. Resnati, E. Riva, T. Perini, E. Ruggieri, U. Orfanelli, F. Paradiso, F. Cremasco, A. Raimondi, E. Pasqualetto, M. Nuvolone, L. Rampoldi, S. Cenci, E. Milan, The interaction of the tumor suppressor FAM46C with p62 and FNDC3 proteins integrates protein and secretory homeostasis. Cell Rep. 32, 108162 (2020).

27. A. B. Herrero, D. Quwaider, L.A. Corchete, M. V. Mateos, R. Garcia-Sanz, N. C. Gutierrez, FAM46C controls antibody production by the polyadenylation of immunoglobulin mRNAs and inhibits cell migration in multiple myeloma. J Cell. Mal. Med. 24, 4171–4182 (2020).

28. O. Gewartowska, G. Aranaz-Novaliches, P. S. Krawczyk, S. Mroczek, M. Kusio-Kobialka, B. Tarkowski, F. Spoutil, O. Benada, O. Kofroiiova, P. Szwedziak, D. Cysewski, J. Gruchota, M. Szpila, A. Chlebowski, R. Sedlacek, J. Prochazka, A. Dziembowski, Cytoplasmic polyadenylation by TENT5A is required for proper bone formation. Cell Rep. 35, 109015 (2021).

29. M. Brouze, A. Czarnocka-Cieciura, 0. Gewartowska, M. Kusio-Kobialka, K. Jachacy, M. Szpila, B. Tarkowski, J. Gruchota, P. Krawczyk, S. Mroczek, E. Borsuk, A. Dziembowski, TENTS-mediated polyadenylation of mRNAs encoding secreted proteins is essential for gametogenesis in mice. Nat. Commun. 15, 5331 (2024).

30. N. Manfrini, M. Mancino, A. Miluzio, S. Oliveto, M. Balestra, P. Calamita, R. Alfieri, R. L. Rossi, M. Sassoe-Pognetto, C. Salio, A. Cuomo, T. Bonaldi, M. Manfredi, E. Marengo, E. Ranzato, S. Martinotti, D. Cittaro, G. Tonon, S. Biffo, FAM46C and FNDC3A are multiple myeloma tumor suppressors that act in concert to impair clearing of protein aggregates and autophagy. Cancer Res. 80, 4693–4706 (2020).

31. S. Yu, V. N. Kim, A tale of non-canonical tails: gene regulation by post-transcriptional RNA tailing. Nat. Rev. Mal. Cell Biol. 21, 542–556 (2020).

32. M. Doyard, S. Bacrot, C. Huber, M. Di Rocco, A. Goldenberg, M. S. Aglan, P. Brunelle, S. Temtamy, C. Michot, G. A. Otaify, C. Baudry, M. Castanet, J. Leroux, J.-P. Bonnefont, A. Munnich, G. Baujat, P. Lapunzina, S. Monnot, V. L. Ruiz-Perez, V. Cormier-Daire, FAM46A mutations are responsible for autosomal recessive osteogenesis imperfecta. J Med. Genet. 55, 278–284 (2018).

33. C. Zheng, Y.-C. Ouyang, B. Jiang, X. Lin, J. Chen, M.-Z. Dong, X. Zhuang, S. Yuan, Q.-Y. Sun, C. Han, Non-canonical RNA polyadenylation polymerase FAM46C is essential for fastening sperm head and flagellum in mice. Biol. Reprod. 100, 1673–1685 (2019).

34. K. D. Boyd, F. M. Ross, B. A. Walker, C. P. Wardell, W. J. Tapper, L. Chiecchio, G. Dagrada, Z. J. Konn, W. M. Gregory, G. H. Jackson, J. A. Child, F. E. Davies, G. J. Morgan, NCRI Haematology Oncology Studies Group, Mapping of chromosome 1p deletions in myeloma identifies FAM46C at lp12 and CDKN2C at lp32.3 as being genes in regions associated with adverse survival. Clin. Cancer Res. 17, 7776–7784 (2011).

35. M. Barbieri, M. Manzoni, S. Fabris, G. Ciceri, K. Todoerti, V. Simeon, P. Musto, A. Cortelezzi, L. Baldini, A. Neri, M. Lionetti, Compendium ofFAM46C gene mutations in plasma cell dyscrasias. Br. J Haematol. 174, 642–645 (2016).

36. T. Liang, X. Ye, Y. Liu, X. Qiu, Z. Li, B. Tian, D. Yan, FAM46B inhibits cell proliferation and cell cycle progression in prostate cancer through ubiquitination of β-catenin. Exp. Mol. Med. 50, 1–12 (2018).

37. Y. X. Zhu, C.-X. Shi, L.A. Bruins, P. Jedlowski, X. Wang, K. M. Kortum, M. Luo, J.M. Ahmann, E. Braggio, A. K. Stewart, Loss of FAM46C promotes cell survival in myeloma. Cancer Res. 77, 4317–4327 (2017).

38. K. Kazazian, Y. Haffani, D. Ng, C. M. M. Lee, W. Johnston, M. Kim, R. Xu, K. Pacholzyk, F. S.-W. Zih, J. Tan, A. Smrke, A. Pollett, H. S.-T. Wu, C. J. Swallow, FAM46C/TENT5C functions as a tumor suppressor through inhibition of Plk4 activity. Commun. Biol. 3,448 (2020).

39. H. Chen, D. Lu, G. Shang, G. Gao, X. Zhang, Structural and functional analyses of the FAM46C/Plk4 complex. Structure 28, 910–921.e4 (2020).

40. Q.-Y. Zhang, X.-Q. Yue, Y.-P. Jiang, T. Han, H.-L. Xin, FAM46C is critical for the anti-proliferation and pro-apoptotic effects of norcantharidin in hepatocellular carcinoma cells. Sci. Rep. 7, 396 (2017).

41. H. Sang, S. Wu, X. Chen, S. Cheng, Q. Li, FAM46B suppresses proliferation, migration and invasion of non-small cell lung cancer via β-catenin/MMP7 signaling. Transl. Cancer Res. 8, 1497–1505 (2019).

42. B. Tajer, J. A. Dutko, S. C. Little, M. C. Mullins, BMP heterodimers signal via distinct type I receptor class functions. Proc. Natl. Acad. Sci. US. A. 118, e2017952118 (2021).

43. J. A. Tucker, K. A. Mintzer, M. C. Mullins, The BMP signaling gradient patterns dorsoventral tissues in a temporally progressive manner along the anteroposterior axis. Dev. Cell 14, 108–119 (2008).

44. M. Hashiguchi, M. C. Mullins, Anteroposterior and dorsoventral patterning are coordinated by an identical patterning clock. Development 140, 1970–1980 (2013).

45. J. S. Park, A. M. Gabel, P. Kassir, L. Kang, P. K. Chowdhary, A. Osei-Ntansah, N. D. Tran, S. Viswanathan, B. Canales, P. Ding, Y.-S. Lee, R. Brewster, N-myc downstream regulated gene 1 (ndrgl) functions as a molecular switch for cellular adaptation to hypoxia. Elife 11 (2022).

46. L. Guglielmi, C. Heliot, S. Kumar, Y. Alexandrov, I. Gori, F. Papaleonidopoulou, C. Barrington, P. East, A. D. Economou, P. M. W. French, J. McGinty, C. S. Hill, Smad4 controls signaling robustness and morphogenesis by differentially contributing to the Nodal and BMP pathways. Nat. Commun. 12, 6374 (2021).

47. H. Greenfeld, J. Lin, M. C. Mullins, The BMP signaling gradient is interpreted through concentration thresholds in dorsal-ventral axial patterning. PLoS Biol. 19, e3001059 (2021).

48. L.A. Passmore, J. Coller, Roles of mRNA poly(A) tails in regulation of eukaryotic gene expression. Nat. Rev. Mol. Cell Biol. 23, 93–106 (2022).

49. K. Xiang, J. Ly, D. P. Bartel, Control ofpoly(A)-tail length and translation in vertebrate oocytes and early embryos. Dev. Cell 59, 1058–1074.el 1 (2024).

50. K. Xiang, D. P. Bartel, The molecular basis of coupling between poly(A)-tail length and translational efficiency. eLife 10 (2021).

51. D. Gaidatzis, L. Burger, M. Florescu, M. B. Stadler, Analysis of intronic and exonic reads in RNA-seq data characterizes transcriptional and post-transcriptional regulation. Nat. Biotechnol. 33, 722–729 (2015).

52. O. Begik, G. Diensthuber, H. Liu, A. Delgado-Tejedor, C. Kontur, A. M. Niazi, E. Valen, A. J. Giraldez, J.-D. Beaudoin, J. S. Mattick, E. M. Novoa, Nano3P-seq: transcriptome-wide analysis of gene expression and tail dynamics using end-capture nanopore cDNA sequencing. Nat. Methods 20, 75–85 (2023).

53. 53. O. Begik, L. P. Pryszcz, A. M. Niazi, E. Valen, E. M. Novoa, Nano3P-seq: charting the coding and noncoding transcriptome at single-molecule resolution. Nat. Protoc. 20, 3607–3628 (2025).

54. S. A. Welsh, A. Gardini, Genomic regulation of transcription and RNA processing by the multitasking Integrator complex. Nat. Rev. Mol. Cell Biol. 24, 204–220 (2023).

55. N. Ma, Y.-K. Wang, S. Xu, Q.-Z. Ni, Q.-W. Zheng, B. Zhu, H.-J. Cao, H. Jiang, F.-K. Zhang, Y.-M. Yuan, E.-B. Zhang, T.-W. Chen, J. Xia, X.-F. Ding, Z.-H. Chen, X.-P. Zhang, K. Wang, S.-Q. Cheng, L. Qiu, Z.-G. Li, Y.-C. Yu, X.-F. Wang, B. Zhou, J.-J. Li, D. Xie, PPDPF alleviates hepatic steatosis through inhibition of mTOR signaling. Nat. Commun. 12, 3059 (2021).

56. T. Watanabe, T. Yamamoto, K. Tsukano, S. Hirano, A. Horikawa, T. Michiue, Fam46a regulates BMP-dependent pre-placodal ectoderm differentiation in Xenopus. Development 145, dev166710 (2018).

57. A. E. Melby, C. Beach, M. Mullins, D. Kimelman, Patterning the early zebrafish by the opposing actions of bozozok and vox/vent. Dev. Biol. 224, 275–285 (2000).

58. Y. Imai, M.A. Gates, A. E. Melby, D. Kimelman, A. F. Schier, W. S. Talbot, The homeobox genes vox and vent are redundant repressors of dorsal fates in zebrafish. Development 128, 2407–2420 (2001).

59. A. Kawahara, T. Wilm, L. Solnica-Krezel, I. B. Dawid, Antagonistic role of vegal and bozozok/dharma homeobox genes in organizer formation. Proc. Natl. Acad. Sci. U. S. A. 97, 12121–12126 (2000).

60. D. Santos, A. Luzio, A. M. Coimbra, Zebrafish sex differentiation and gonad development: A review on the impact of environmental factors. Aquat. Toxicol. 191, 141–163 (2017).

61. N. D. Meeker, S. A. Hutchinson, L. Ho, N. S. Trede, Method for isolation of PCR-ready genomic DNA from zebrafish tissues. Biotechniques 43,610,612,614 (2007).

62. C. Gerri, R. Marin-Juez, M. Marass, A. Marks, H.-M. Maischein, D. Y. R. Stainier, Hif-la regulates macrophage-endothelial interactions during blood vessel development in zebrafish. Nat. Commun. 8, 15492 (2017).

63. J. Zinski, F. Tuazon, Y. Huang, M. Mullins, D. Umulis, Imaging and quantification of P-Smadl/5 in zebrafish blastula and gastrula embryos. Methods Mol. Biol. 1891, 135–154 (2019).

64. J. Zinski, Y. Bu, X. Wang, W. Dou, D. Umulis, M. C. Mullins, Systems biology derived source-sink mechanism ofBMP gradient formation. Elife 6 (2017).

65. S. H. Colbert, A. F. Mack, R. D. Fernald, A novel, rapid flat-mounting technique for visualizing antibody labeling in the retina. J Neurosci. Methods 62, 179–183 (1995).

66. D. E. Wagner, C. Weinreb, Z. M. Collins, J. A. Briggs, S. G. Megason, A. M. Klein, Single-cell mapping of gene expression landscapes and lineage in the zebrafish embryo. Science 360, 981–987 (2018).

67. N. D. Lawson, R. Li, M. Shin, A. Grosse, O. Yukselen, O. A. Stone, A. Kucukural, L. Zhu, An improved zebrafish transcriptome annotation for sensitive and comprehensive detection of cell type-specific genes. Elife 9 (2020).

68. F. A. Wolf, P. Angerer, F. J. Theis, SCANPY: large-scale single-cell gene expression data analysis. Genome Biol. 19, 15 (2018).

69. S. L. Wolock, R. Lopez, A. M. Klein, Scrublet: Computational identification of cell Doublets in Single-cell transcriptomic data. Cell Syst. 8, 281–291.e9 (2019).

70. C. Dominguez Conde, C. Xu, L.B. Jarvis, D. B. Rainbow, S. B. Wells, T. Gomes, S. K. Howlett, O. Suchanek, K. Polanski, H. W. King, L. Mamanova, N. Huang, P.A. Szabo, L. Richardson, L. Bolt, E. S. Fasouli, K. T. Mahbubani, M. Prete, L. Tuck, N. Richoz, Z. K. Tuong, L. Campos, H. S. Mousa, E. J. Needham, S. Pritchard, T. Li, R. Elmentaite, J. Park, E. Rahmani, D. Chen, D. K. Menon, O. A. Bayraktar, L. K. James, K. B. Meyer, N. Yosef, M. R. Clatworthy, P.A. Sims, D. L. Farber, K. Saeb-Parsy, J. L. Jones, S. A. Teichmann, Cross-tissue immune cell analysis reveals tissue-specific features in humans. Science 376, eabl5197 (2022).

71. A. M. Klein, L. Mazutis, I. Akartuna, N. Tallapragada, A. Veres, V. Li, L. Peshkin, D. A. Weitz, M. W. Kirschner, Droplet barcoding for single-cell transcriptomics applied to embryonic stem cells. Cell 161, 1187–1201 (2015).

72. J.B. Kang, A. Nathan, K. Weinand, F. Zhang, N. Millard, L. Rumker, D. B. Moody, I. Korsunsky, S. Raychaudhuri, Efficient and precise single-cell reference atlas mapping with Symphony. Nat. Commun. 12, 5890 (2021).

73. I. Korsunsky, N. Millard, J. Fan, K. Slowikowski, F. Zhang, K. Wei, Y. Baglaenko, M. Brenner, P.-R. Loh, S. Raychaudhuri, Fast, sensitive and accurate integration of single-cell data with Harmony. Nat. Methods 16, 1289–1296 (2019).

74. 74. L. Mcinnes, J. Healy, J. Melville, UMAP: Uniform Manifold Approximation and Projection for Dimension Reduction, arXiv [stat.ML] (2018). http://arxiv.org/abs/1802.03426.

75. B. Muzellec, M. Telenczuk, V. Cabeli, M. Andreux, PyDESeq2: a python package for bulk RNA-seq differential expression analysis. Bioinformatics 39, btad547 (2023).

76. P. Di Tommaso, M. Chatzou, E.W. Floden, P. P. Barja, E. Palumbo, C. Notredame, Nextflow enables reproducible computational workflows. Nat. Biotechnol. 35, 316–319 (2017).

77. P.A. Ewels, A. Peltzer, S. Fillinger, H. Patel, J. Alneberg, A. Wilm, M. U. Garcia, P. Di Tommaso, S. Nahnsen, The nf-core framework for community-curated bioinformatics pipelines. Nat. Biotechnol. 38, 276–278 (2020).

78. A. Dobin, C. A. Davis, F. Schlesinger, J. Drenkow, C. Zaleski, S. Jha, P. Batut, M. Chaisson, T. R. Gingeras, STAR: ultrafast universal RNA-seq aligner. Bioinformatics 29, 15–21 (2013).

79. R. Patro, G. Duggal, M. I. Love, R. A. Irizarry, C. Kingsford, Salmon provides fast and bias-aware quantification of transcript expression. Nat. Methods 14, 417–419 (2017).

80. M. I. Love, W. Huber, S. Anders, Moderated estimation of fold change and dispersion for RNA-seq data with DESeq2. Genome Biol. 15, 550 (2014).

81. K. Blighe, S. Rana, M. Lewis, EnhancedVolcano: Publication-ready volcano plots with enhanced colouring and labeling. R package version (2019).

82. G. Yu, L.-G. Wang, Y. Han, Q.-Y. He, clusterProfiler: an R package for comparing biological themes among gene clusters. OMICS 16, 284–287 (2012).

83. F. Krueger, Trim Galore!: A wrapper around Cutadapt and FastQC to consistently apply adapter and quality trimming to FastQ files, with extra functionality for RRBS data. Babraham Institute (2015).

84. H. Li, R. Durbin, Fast and accurate short read alignment with Burrows-Wheeler transform. Bioinformatics 25, 1754–1760 (2009).

85. Y. Zhang, T. Liu, C. A. Meyer, J. Eeckhoute, D. S. Johnson, B. E. Bernstein, C. Nusbaum, R. M. Myers, M. Brown, W. Li, X. S. Liu, Model-based analysis of ChIP-Seq (MACS). Genome Biol. 9, R137 (2008).

86. C. S. Ross-Innes, R. Stark, A. E. Teschendorff, K. A. Holmes, H. R. Ali, M. J. Dunning, G. D. Brown, O. Gojis, I. O. Ellis, A. R. Green, S. Ali, S.-F. Chin, C. Palmieri, C. Caldas, J. S. Carroll, Differential oestrogen receptor binding is associated with clinical outcome in breast cancer. Nature 481, 389–393 (2012).

87. P. P. Singh, G. A. Reeves, K. Contrepois, K. Papsdorf, J. W. Miklas, M. Ellenberger, C.-K. Hu, M. P. Snyder, A. Brunet, Evolution of diapause in the African turquoise killifish by remodeling the ancient gene regulatory landscape. Cell 187, 3338–3356.e30 (2024).

88. G. Yu, L.-G. Wang, Q.-Y. He, ChIPseeker: an R/Bioconductor package for ChIP peak annotation, comparison and visualization. Bioinformatics 31, 2382–2383 (2015).

89. I. Seiliez, B. Thisse, C. Thisse, FoxA3 and goosecoid promote anterior neural fate through inhibition of Wnt8a activity before the onset of gastrulation. Dev. Biol. 290, 152–163 (2006).

90. I. A. Swinburne, K. R. Mosaliganti, A. A. Green, S. G. Megason, Improved Long-Term Imaging of Embryos with Genetically Encoded a-Bungarotoxin. PLoS One 10, e0134005 (2015).

91. H. Gatzke, M. Kilisch, M. Martinez-Carranza, S. Sograte-Idrissi, A. Rajavel, T. Schlichthaerle, N. Engels, R. Jungmann, P. Stenmark, F. Opazo, S. Frey, The ALFA-tag is a highly versatile tool for nanobody-based bioscience applications. Nat. Commun. 10, 4403 (2019).

92. C. W. Boswell, C. Hoppe, A. Sherrard, L. Miao, M. L. Kojima, P. Martino, N. Zhao, T. J. Stasevich, S. Nicoli, A. J. Giraldez, Genetically encoded affinity reagents are a toolkit for visualizing and manipulating endogenous protein function in vivo. Nat. Commun. 16, 5503 (2025).

93. A. Brouze, P. S. Krawczyk, A. Dziembowski, S. Mroczek, Measuring the tail: Methods for poly(A) tail profiling. Wiley Interdiscip. Rev. RNA 14, el 737 (2023).

94. H. Chang, J. Lim, M. Ha, V. N. Kim, TAIL-seq: genome-wide determination ofpoly(A) tail length and 3’ end modifications. Mal. Cell 53, 1044–1052 (2014).

